# Reversal of lineage plasticity in RB1/TP53-deleted prostate cancer through FGFR and Janus kinase inhibition

**DOI:** 10.1101/2021.11.01.466615

**Authors:** Wouter R. Karthaus, Joseph Chan, Manu Setty, Jillian R. Love, Samir Zaidi, Zi-ning Choo, Sitara Persad, Justin LaClair, Kayla E Lawrence, Ojasvi Chaudhary, Tianhao Xu, Ignas Masilionis, Linas Mazutis, Ronan Chaligne, Dana Pe’er, Charles L Sawyers

## Abstract

The inherent plasticity of tumor cells provides a mechanism of resistance to many molecularly targeted therapies, exemplified by adeno-to-neuroendocrine lineage transitions seen in prostate and lung cancer. Here we investigate the root cause of this lineage plasticity in a primary murine prostate organoid model that mirrors the lineage transition seen in patients. These cells lose luminal identity within weeks following deletion of *Trp53* and *Rb1*, ultimately acquiring an Ar-negative, Syp+ phenotype after orthotopic *in vivo* transplantation. Single-cell transcriptomic analysis revealed progressive mixing of luminal-basal lineage features after tumor suppressor gene deletion, accompanied by activation of Jak/Stat and Fgfr pathway signaling and interferon-a and -g gene expression programs prior to any morphologic changes. Genetic or pharmacologic inhibition of Jak1/2 in combination with FGFR blockade restored luminal differentiation and sensitivity to antiandrogen therapy in models with residual AR expression. Collectively, we show lineage plasticity initiates quickly as a largely cell-autonomous process and, through newly developed computational approaches, identify a pharmacological strategy that restores lineage identity using clinical grade inhibitors.

## Main

Although resistance to molecularly targeted therapies is often due to mutations in the drug target, there is growing recognition of other modes of tumor escape, particularly with next-generation inhibitors designed to circumvent target-based resistance mechanisms. For example, in EGFR- and KRAS-mutant lung cancer and in metastatic prostate cancer, tumor cells can undergo a lineage transition from adenocarcinoma to squamous or neuroendocrine histology following treatment with the respective inhibitors (*1-6*). Similarly, BRAF-mutant melanomas can transition from a MITF+ differentiated melanocyte phenotype to a more mesenchymal AXL+ cell state or neural crest stem-like state in response to BRAF inhibition (*7-9*). Transcriptomic analyses of these changing cell states share features of normal developmental, regenerative and stem cell signatures, suggesting that tumor cells co-opt these gene expression programs to escape lineage-dependent therapies (*10*). Importantly, these plasticity mechanisms are not associated with new genomic alterations and are potentially reversible (*11, 12*).

The increasing prevalence of this mode of tumor escape has sparked numerous investigations into the underlying mechanism. Analysis of patient specimens, particularly at the single-cell level, has been particularly instructive in defining the complexity of the problem, revealing remarkable heterogeneity within and across patients, as exemplified in our accompanying manuscript and by others (*13, 14*). Because these studies typically capture a single snapshot during tumor evolution, it is difficult to discern precisely how plasticity arises. Furthermore, the complexity of genomic alterations in advanced cancers makes it challenging to distinguish between drivers of oncogenicity versus drivers of plasticity-associated drug resistance. To complement these patient-focused studies, we and others have identified several genetic perturbations that can induce plasticity and drug resistance in prostate cancer models, such as loss of tumor suppressor genes (TP53, RB1, PTEN) and activation of transcription factors (MYC, NMYC, SOX2, BRN2) in various combinations (*11, 15-17*). In some cases, these same perturbations can also elicit plasticity phenotypes in lung, providing further evidence for the developmental conservation of these programs across endoderm-derived epithelial tissues(*5, 15*).

We previously reported conditions for *in vitro* propagation and genetic manipulation of primary prostate organoids under defined, serum-free growth conditions (*18*). Because co-deletion of *Trp53* and *Rb1* in the mouse prostate has been shown to give rise to carcinomas displaying both luminal epithelial and neuroendocrine differentiation(*19*), we asked if a similar phenotype might be seen *ex vivo* by infection of *Trp53^loxP/loxP^, Rb1 ^loxP/loxP^* organoids with Cre-expressing lentivirus. The presence of a mTmG reporter cassette allowed easy identification and tracking of *Trp53^-/-^*; *Rb^-/-^* cells due to the red-to-green color change initiated by Cre expression (**Movie S1**). In contrast to the cystic appearance of uninfected organoids, those with *Trp53/Rb1* co-deletion lost their basal/luminal polarity and cystic lumens by ∼4 weeks (hereafter called hyperplastic), followed by the appearance of migratory cells and cellular protrusions outside the normally sharp exterior organoid borders at ∼8-10 weeks (hereafter called “slithering,” based on similarity to the migratory phenotype reported for lung neuroepithelial bodies) (*20*)(**Fig 1A-B, Fig S1A, Fig S2, Movie S2**). This phenotype required deletion of both tumor suppressors and was also seen after CRISPR/Cas9-mediated co-deletion of *Trp53* and *Rb1* in wild-type organoids (**Fig S1B**).

**Figure 1.**
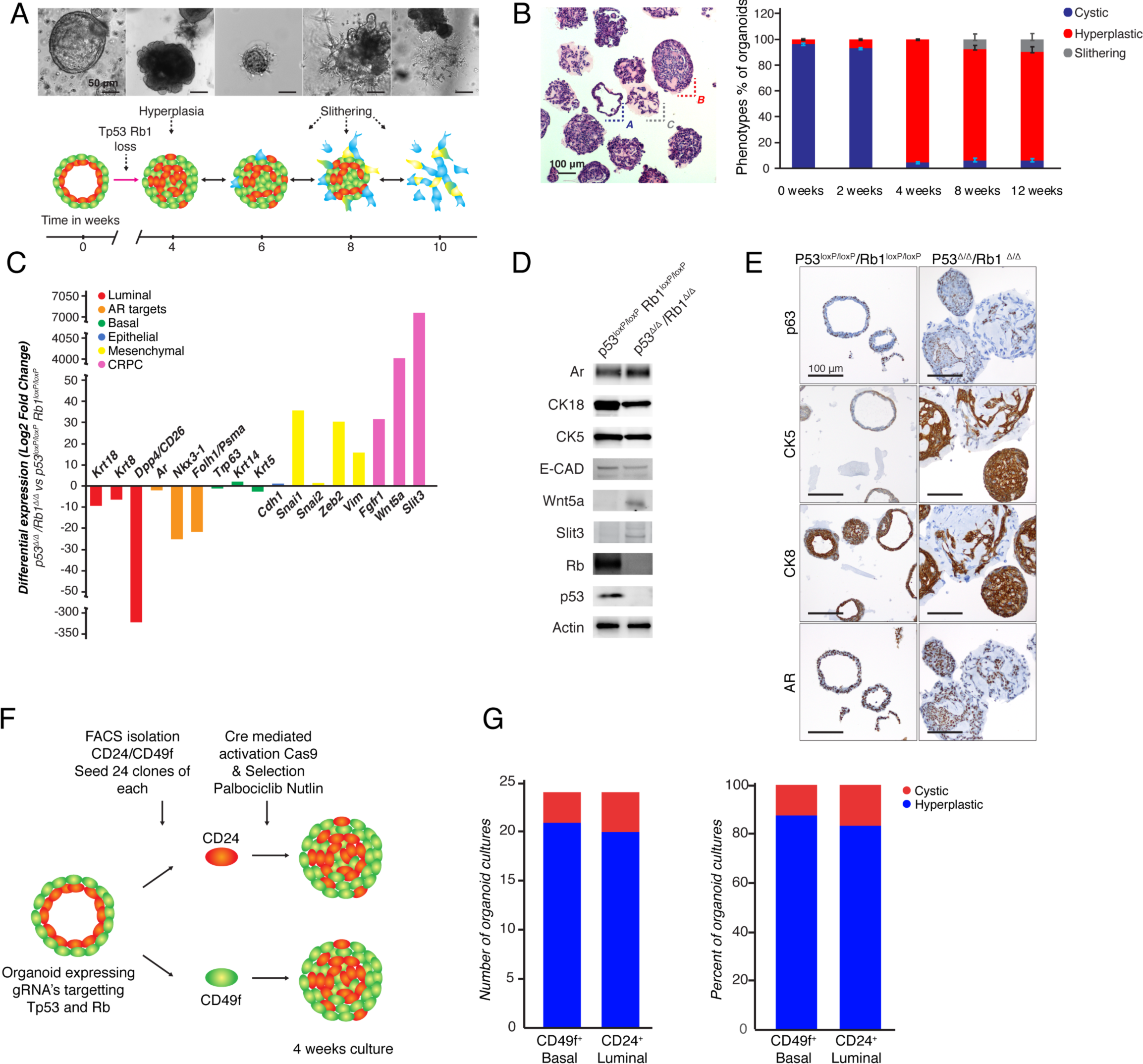
Establishment of an organoid model of spontaneous lineage plasticity: **A)** *Upper:* Representative brightfield pictures of the various organoid phenotypes prior to and 4,6,8 and 10 weeks post *Tp53* and *Rb1* loss. *Lower:* Schematic representation of lineage plasticity development in *Tp53^Δ/Δ^ Rb1^Δ/Δ^* organoids. Scale bar represents 50 μm. **B)** *Left:* Representative H&E staining of LentiCre *Tp53^Δ/Δ^ Rb1^Δ/Δ^* organoids ∼10 weeks post deletion with cystic (A, blue), Hyperplastic (B, red) and slithering (C, silver) phenotypes. *Right:* Bargraph with percentage of organoids in given phenotypes during the time course. Scale bar represents 100 μm. **C)** Bargraph of differentially expressed genes between ∼10 weeks post deletion (LentiCre *Tp53^Δ/Δ^ Rb1^Δ/Δ^* organoids) and wild-type (*Tp53^loxP/loxP^ Rb1^loxP/loxP^* organoids) in bulk RNA-seq. DESeq2, FDR ≤ 0.05, Log2 fold change (for full parameters see **Methods**). Genes are colored by association to lineage (Luminal (Red), AR target (Orange), Basal (Green), Epithelial (Blue), Mesenchymal (Yellow), and CRPC (Magenta). Neuroendocrine marker genes were lowly expressed without major difference (*Cghb, Syp*) or not detected (*Cgha, NeuroD1, Ascl1*) in RNA-seq data (see supplementary Table 1). **D)** Westernblot verification of differentially expressed genes identified in RNA-seq data in wild-type (*Tp53^loxP/loxP^ Rb1^loxP/loxP^* organoids, left) and ∼10 weeks post deletion (LentiCre *Tp53^Δ/Δ^ Rb1^Δ/Δ^* organoids, right). Proteins as marked. Actin was used as a loading control. **E)** Representative IHC of basal markers (p63, Ck5), luminal marker (Ck8) and Ar in wild-type (*Tp53^loxP/loxP^ Rb1^loxP/loxP^* organoids, left) and ∼10 weeks post deletion (LentiCre *Tp53^Δ/Δ^ Rb1^Δ/Δ^* organoids, right). Scale bars represent 100 μm. **F)** Schematic representation of single cell cloning experiment of Basal (CD49f^+^) and Luminal (CD24^+^) cells. Organoids harboring a Cre recombinase inducible Cas9 (Rosa26 lsl-Cas9 (*32*) were transduced with guide RNA’s targeting *Tp53* and *Rb1* (LentiGuide Puro, guide sequences **Table S**). Basal and luminal cells were sorted based on CD49f^+^ (Basal) and CD24^+^ (Luminal) expression and subsequently Cas9 was activated by transduction with adenovirus expressing Cre. Approximately 1000 single cells were seeded and grown out. Nutlin-3 and Palbociclib were added 3 days post Cas9 activation to select for *Tp53* and *Rb1* mutant cells. 24 basal and 24 luminal clones were randomly chosen and expanded for 4 weeks. **G)** Quantification whole culture phenotypes (Cystic or Hyperplastic) of single cell derived (CD49f^+^ or CD24^+^ FACS sorted) organoids after 4 weeks of culture. Left bargraph absolute number of cultures, right bargraph percentage of cultures in cystic or hyperplastic state.

To further characterize the changes induced by *Trp53/Rb1* co-deletion, we performed bulk RNA-seq, western blot analysis and *in situ* immunofluorescence (IF) or immunohistochemistry (IHC) for various lineage markers. The most dramatic finding was reduced expression of luminal lineage genes such as *Nkx3.1, Folh1* (Psma), *Dpp4* (CD26), *Krt18* and *Krt8* and increased expression of mesenchymal genes such as *Snai1, Zeb2, Vim* (**Fig 1C-D, S1C-D, Table S1**). Other notable gene expression changes include *Wnt5a* and the axon guidance ligand *Slit3*, both of which can promote migratory behavior, and *Fgfr1* which has been implicated in epithelial-mesenchymal transition (EMT) associated drug resistance in prostate cancer as well as EGFR-mutant lung cancer (*21, 22*). However, unlike the Syp+ neuroendocrine cancer seen in the *Trp53^-/-^;Rb1^-/-^* mouse prostate cancer model, we observed only modest levels of *Syp* and *Chgb* and failed to detect expression of *Chga, Ascl1* or *Neurod1*, even in *Trp53^-/-^;Rb1^-/-^* organoids passaged for >12 months.

We postulated the lack of Syp or other neuroendocrine lineage gene expression might be due to absence of critical prostate microenvironmental factors in organoid culture media, and therefore asked if the organoid phenotype evolved further after orthotopic (OT) implantation of *Trp53^-/-^/Rb1^-/-^* organoids into *Trp53^+/+^*; *Rb1^+/+^* hosts. After confirming highly efficient engraftment of hyperplastic/slithering stage organoids at 4 weeks in a pilot experiment (by detection of GFP+ cells in the dorsal lobe of hormonally intact *NSG* mice), we scored lineage phenotype and tumorigenicity, as well as the effect of androgen blockade, in a larger cohort by treating half the mice with castration and daily enzalutamide (Enz) 4 weeks after OT injection. In the hormonally intact mice, engrafted cells remained Ar+, Syp- (as in organoid culture); however, large foci of Ar-, Syp+ engrafted cells were seen in all mice in the castration + Enz group, indicating progression to a neuroendocrine-like lineage (**Fig S3A-B**). This progression to an Ar-, Syp+ state following androgen receptor signaling inhibition (ARSI) was confirmed across four independent *Trp53/Rb1* co-deletion experiments (**Fig S3C**). To determine if this effect of *in vivo* androgen blockade on lineage plasticity is also observed in culture, we returned to the organoid model and compared the effect of dihydrotestosterone (DHT) versus Enz on the slithering phenotype following *Trp53/Rb1* co-deletion. The percentage of organoids scored as slithering increased from ∼10% to >30% (∼3-fold) after 7 days of Enz treatment **(Fig S3D**). Thus, sustained Ar signaling plays a role in preserving luminal lineage identity, a conclusion further supported by recent data from a human patient-derived xenograft model (*23*).

The above experiments demonstrate that lineage plasticity can be initiated by co-deletion of *Trp53* and Rb1 in normal mouse prostate organoids grown in bulk culture, which includes basal and luminal cells. To determine if acquisition of plasticity is restricted to either of these populations, we isolated basal (CD49f+) and luminal (CD24+) cells by FACS prior to lentiviral Cre infection and found that both basal and luminal cells can give rise to plasticity in response to co-deletion of *Trp53* and *Rb1* (**Fig 1F-G, S1E-F**).

Having developed a predictable model in which lineage plasticity initiates and evolves, we wanted to characterize the emerging plasticity and elucidate the molecular mechanisms through which this plasticity is induced following co-deletion of *Trp53* and *Rb1*. To characterize the transcriptional cell states and gene programs that emerge over time, we collected single-cell RNA-sequencing (scRNA-seq) data, beginning 2 weeks after *Trp53/Rb1* co-deletion, prior to any morphologic evidence of plasticity, then at 4 and 8 weeks (**Fig 2A; Table S2**). These experiments were done in bulk-derived organoids in the presence of either DHT or Enz based on the effects of androgen blockade on plasticity described above (**Fig S3**).

**Figure 2:**
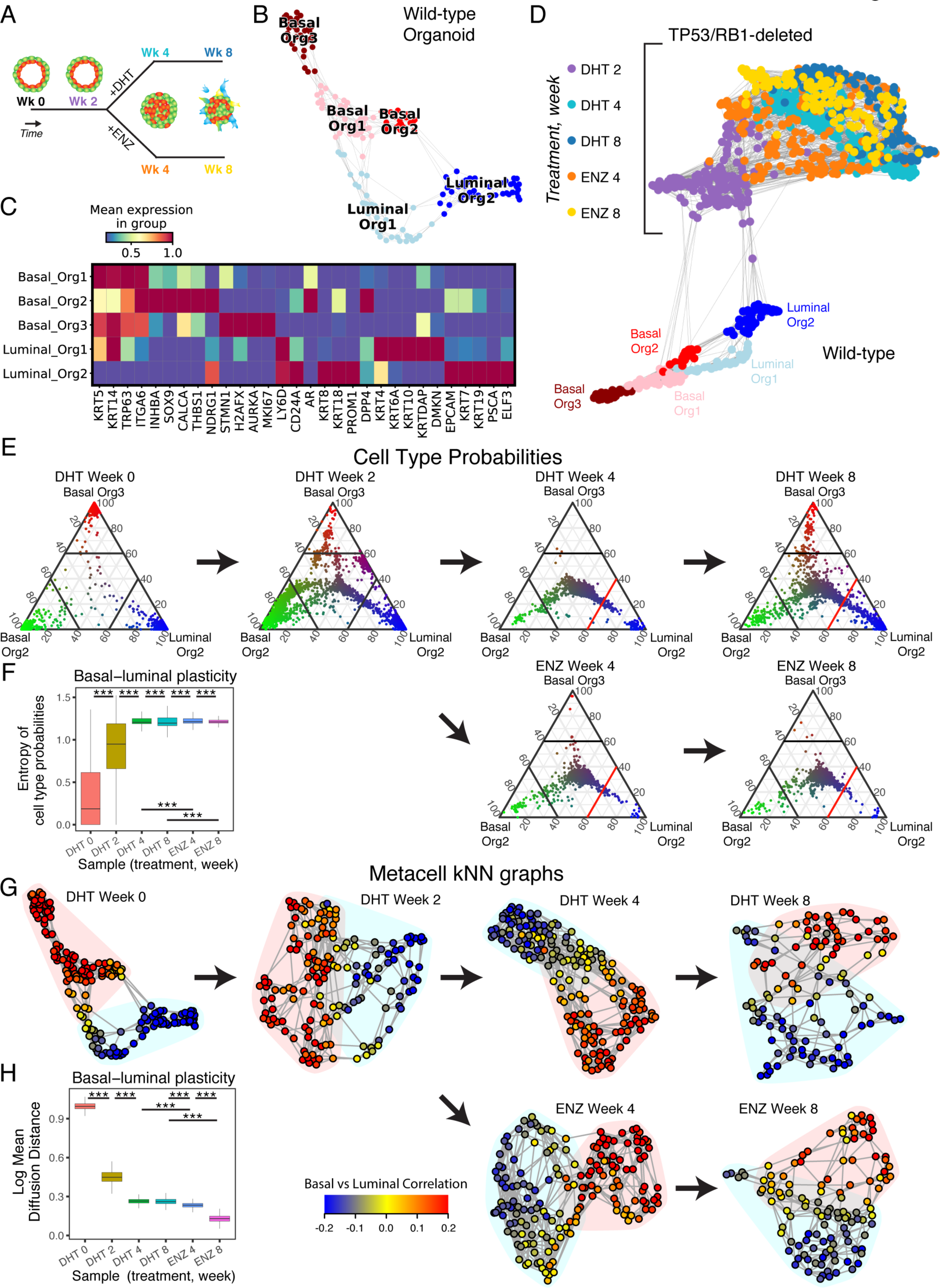
Single-cell sequencing reveals increased basal-luminal plasticity in the prostate organoid following *RB1/TP53* deletion. **A)** Experimental design of scRNA-seq time-course sequenced at weeks 0, 2, 4, and 8, with weeks 4 and 8 subjected to either enzalutamide (ENZ) treatment or dihydrotestosterone (DHT) control. Each sample then undergoes enzymatic digestion and subsequent scRNA-seq. **B)** Force-directed layout (FDL) of wild-type basal and luminal SEACell metacells, labeled by wild-type subpopulations determined by Phenograph clusters. Edges between metacells indicate the k-nearest neighbors (k = 6) (see **Methods**). **C)** Heatmap of mean log2(X+1) expression of DEGs per basal and luminal subpopulation in the wild-type organoid. **D)** FDL (k=10) of metacells in the prostate organoid before and after *RB1/TP53* deletion, annotated by either wild-type subpopulations or by time and treatment following mutation. **E)** Ternary plots of cell-type classification (based on Markov absorption probabilities) for single cells in the organoid before and after mutation. Each ternary plot represents a timepoint/condition. Each dot is a single cell, plotted using 3 coordinates representing the classification probabilities of the 3 most frequent basal and luminal phenotypes in the wild-type organoid: Basal Org2 (green), Basal Org3 (red), and Luminal Org2 (blue). Color intensity of each dot corresponds to the certainty of cell-type classification, calculated by entropy of their classification probabilities (see **Methods**). Higher entropy indicates greater uncertainty of cell type assignment, increased basal-luminal mixing, a proxy for increased plasticity. The red line at 60% probability of Luminal Org2 classification at weeks 4 and 8 highlights the decreased probability for Luminal Org2 in Enz-treated samples compared to DHT control. **F)** Boxplot of entropy of cell-type classification probabilities for single cells per timepoint in (**G**) as a measure of basal-luminal mixing and plasticity over time and treatment (Wilcoxon rank-sum test). p-values: ***<0.001 **G)** FDL of metacells within each timepoint/treatment. Nodes represent metacells, and edges represent k-nearest neighbors (k = 3% of the number of metacells per timepoint). Metacell nodes are colored based on correlation to bulk-sorted basal CD49f+ and luminal CD24+ cells. Each FDL is divided into two predominantly basal vs luminal partitions, annotated by pale red (basal) and pale blue (luminal) polygons, determined by attributed stochastic block model (aSBM) (**see Methods**). **H)** Boxplot of log mean diffusion distances between basal and luminal partitions for each FDL in (**G**), as an inverse measure of plasticity over time and treatment (Wilcoxon rank-sum test, see **Methods**). p-values: ***<0.001

We observed great transcriptional heterogeneity both within and between each time point/ condition. To catalog systematically the observed cell states that occurred in our prostate system, we started with the wildtype phenotypic landscape then followed how it remodels following perturbation. To gain a concise description of the observed phenotypic diversity, we used our SEACell algorithm (see methods) to calculate metacells separately within each sample timepoint, then ensured that the resulting metacells were robust (**Fig S4**). The metacell approach(*24*) creates a set of distinct, highly granular cell states by aggregating similar cells to generate a robust, full transcriptomic quantification for each cell state, thus mitigating noise and data sparsity in scRNA-seq. An added benefit of metacells is enumeration of observed cell states that are more amenable for comparison (see **Fig S5** for workflow of metacell versus single-cell analyses).

We first focused on the cell states present in the wild-type organoid. Applying Phenograph clustering (*25*), we identified 5 distinct subpopulations, which we classified as luminal (L-Org1, L-Org2) and as basal (B-Org 1, B-Org 2, B-Org 3) based on published transcriptomes of luminal and basal cells (**see Methods**)(**Fig 2B-C, S6A-B, Table S3-8**). L-Org1 cells share expression of stem-like genes (e.g. *Ly6d*) with L-Org2 cells but also express some basal genes (e.g. *Krt14*), perhaps indicative of a bi-potential population enriched in organoid culture. L-Org2 cells resemble stem-like (Ly6d+, Psca+) luminal cells seen *in vivo* (called L2 in Karthaus et al, LumP in Crowley et al) and *Prom1*, which marks secretory luminal cells seen *in vivo* (*26, 27*). B-Org1 cells express canonical basal markers (*Krt5*+, *Krt14+, Tp63*+), B-Org2 cells express secretory proteins such calcitonin and inhibin A, whereas B-Org3 are largely distinguished by expression of proliferation markers (*Mki67, Aurka*).

Analyzing the entire dataset, we found no overlap between the cell states present in the wild-type organoid and the mutant cell states. We visualized all metacells from the entire time-course following *Trp53/Rb1* co-deletion using a force-directed layout (FDL). The metacells organized along the FDL based on time point, with a clearly distinct gap between the wild-type and 2-week timepoint. Strikingly, while the wild-type metacells organized into clear subpopulations, the mutant metacells formed a seemingly disorganized cloud of cell states lacking clear distinction from one another (**Fig 2D**). In a manner similar to the wild-type metacells, we used gene signatures derived from sorted luminal and basal cells to assign luminal versus basal identity to the mutant cells. While we observe clear basal and luminal phenotypes among some mutant metacells, the distinction is far less clear for the majority of the mutant metacells **(Fig S6C**), which express genes associated with both luminal (e.g., *KRT8*) and basal (e.g., *KRT5*) identities (**Fig S6D**). To ensure that the mixed basal-luminal phenotype is not a product of the metacell approach, we repeated the analysis at the level of individual cells and observed similar lineage mixing (**Fig S6E, see Methods**).

To explore and quantify the mixing of basal/luminal identity further, we deconvolved cell phenotype using a Markov-absorption classification approach(*25*) (**see Methods**) which calculates the probability of associating with each of the wild-type sub-populations based on transcriptomic similarity. The key advantage to this graph-based approach is that it scores similarity based on the key axes of gene variation along the phenotypic manifold, giving more weight to the genes that participate in a multitude of intermediate states. As a result, each cell is assigned a vector of classification probabilities for each wild-type subpopulation. While some cells are clearly associated with one subpopulation, others have an uncertain classification indicating basal-luminal mixing. To quantify this mixing, we calculated the entropy of these cell type probabilities and treat this entropy as a proxy measure of plasticity.

To visualize the basal luminal mixing, we plot the classification probabilities for the 3 predominant basal/luminal phenotypes (B-Org2, B-Org3, L-Org2) (**Fig S6F**) as a ternary plot, stratified by sample timepoint (**Fig 2E, Methods**). As expected, the undeleted cells favor a single assignment to their respective populations, but as time progresses, cell type probabilities become less committed to a single basal or luminal phenotype, eventually converging towards maximal uncertainty at the center of the ternary plot. Interestingly, cells from the Enz-treated samples display even fewer cells favoring the L-Org2 phenotype, consistent with our early data showing that Ar inhibition accelerates loss of luminal identity. Using our entropy-based plasticity score, we observed a time-dependent increase following *Trp53/Rb1* co-deletion (**Fig 2F**) that was robust across parameters (**Fig S6G**). To demonstrate robustness further, we repeated the analysis using an alternative approach dependent only on gene signatures. By calculating a normalized average Z-score of DEGs for the undeleted (wild-type) basal and luminal subpopulations for each single cell (**see Methods**), we again observe increased basal-luminal mixing over time, with a phenotypic shift away from luminal phenotype with Enz treatment and an increase in entropy as a measure of plasticity over time (**Fig S6H-I**).

Having observed an increasing fraction of mixed-lineage cells over time, we sought to better characterize the mixed lineage cell-states. At each timepoint, metacells represent distinct cell states, and we visualize their structure using an FDL on the nearest-neighbor graph (**Fig 2G**). The wild-type metacells clearly partition into a luminal and a basal component with limited paths between them. At two weeks, while the organoids are still morphologically intact, we observe a diversity of distinct intermediate states that mediate an increasing number of connections between the basal and luminal components, suggesting multiple potential paths of transdifferentiation. Over time and with Enz treatment, the diversity of intermediate states and the connectivity between basal and luminal components continue to increase, which we quantify using diffusion distance as an inverse proxy measure of plasticity (**Fig 2H, S6J**).

In summary, our single-cell analyses identify transcriptional changes associated with lineage plasticity early within 2 weeks, well in advance of the morphologic changes that evolve over 4-8 weeks. Furthermore, we quantify an increase in both the fraction of mixed lineage cells and diversity of mixed lineage cell-states over both time and treatment. While lineage tracing is required to assess plasticity conclusively, our quantitative measures of entropy and diffusion distance consistently detect increasing plasticity over time, accelerated by drugs that inhibit Ar pathway signaling.

We next sought to elucidate the candidate pathways that increase plasticity in the mutant organoid. We leveraged our measure of plasticity that increases over time yet still displays a wide range within each timepoint/condition. Thus, for each cell, we correlated our measure of plasticity with gene programs using GSEA (**Table S9-10;** see **Methods**). While correlation does not imply causation, we reasoned that gene programs responsible for plasticity would: (1) correlate to our plasticity measure and (2) activate early in the time course when plasticity begins to arise even before morphological changes take place (week 2 following *Trp53/Rb1* deletion). The top candidate pathways were SMAD2 (EMT), LIF signaling, JAK-STAT and FGFR signaling (**Fig 3A-B, S7A-C; Table S11**). In addition to the expected EMT signature (SMAD2) based on our earlier characterization of bulk organoid cultures (**Fig 1D**), we were surprised to find gene sets consistent with inflammation (LIF signaling, JAK-STAT). Indeed, our entropy-based score was tightly correlated with JAK/STAT activity in each cell (**Fig 3C**). Additionally, these cells also display an interferon response signature (**Fig 3B**). We confirmed JAK/STAT signaling at the protein level by western blot analysis showing elevated phospho-Stat1 (pStat1) and phospho-Stat3 following Trp53/Rb1 deletion (**Fig 3D**).

**Figure 3:**
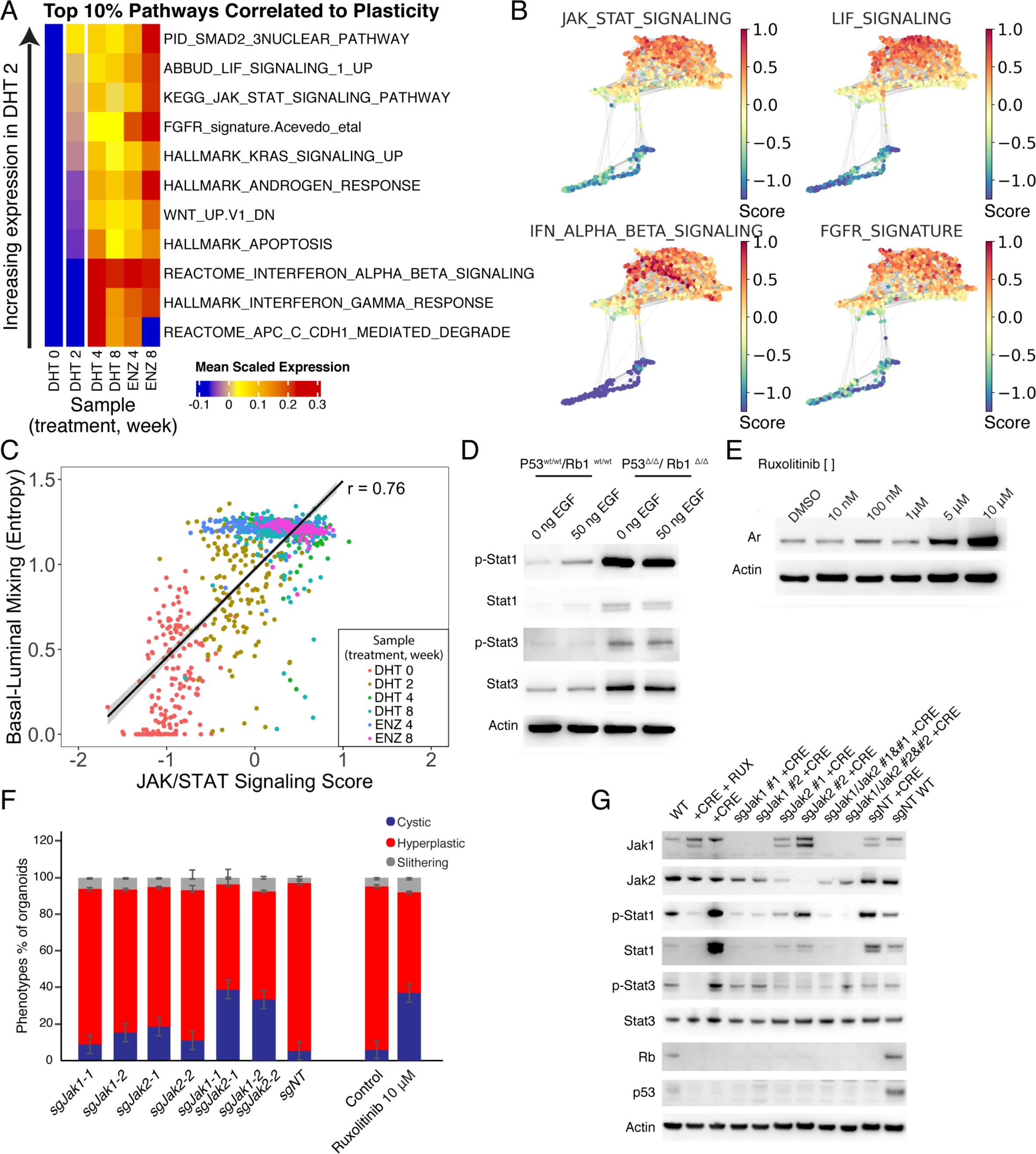
JAK/STAT signaling drives increased plasticity in the *Rb1/Tp53*-deleted prostate organoid. **A)** Heatmap of select pathways from the top 10% correlated to plasticity across timepoints (see **Figure S5C,** Table S11). Each pathway score is measured as the average Z-score of gene expression in each pathway among SEACells. Rows (pathways) are ordered by increasing pathway score from bottom to top in the early timepoint at week 2, with JAK/STAT and LIF signaling scoring the highest (see **Methods**). **B)**, FDL of SEACells in the prostate organoid cells before and after *RB1/TP53* deletion (same layout as **Fig 2D**), scored by key pathways enriched in the mutant setting. Scores are calculated by average Z-score of the leading-edge gene subsets calculated by GSEA for the following pathways from MSigDB: HALLMARK_IL6_JAK_STAT3_SIGNALING, ABBUD_LIF_SIGNALING_1_UP, REACTOME_INTERFERON_ALPHA_BETA_SIGNALING, and a curated signature of FGFR signaling in prostate adenocarcinoma based on Acevedo, et al.(*22*) combined with FGFR1-3 and FGF ligands (see **Methods**). **C)** Scatterplot of JAK/STAT signaling score vs plasticity as measured by average entropy of cell-type classification probabilities per metacell. JAK/STAT score is measured by average Z-score of the leading-edge gene subset of KEGG_JAK_STAT_SIGNALING_PATHWAY and HALLMARK_IL6_JAK_STAT3_SIGNALING. For visualization, the entropy of classification probabilities was first calculated for each single cell and then averaged per metacell. A linear fit is shown with Pearson’s correlation of 0.74. **D)** Westernblot of Stat1 and phosphorylated Stat1 (p-Stat1) in LentiCre *Tp53^Δ/Δ^ Rb1^Δ/Δ^* organoids cultured with 50 ng/ml or 0 ng/ml EGF for 7 days. Actin was used a loading control. **E)** Westernblot of AR in 21-day Ruxolitinib treated (0, 10 nM, 100 nM, 5 μM, 10 μM, 10 μM) organoids **F)** Bargraph with percentage of cystic (Blue), Hyperplastic (Red) and slithering (Silver) phenotypes in LentiCre *Tp53^Δ/Δ^ Rb1^Δ/Δ^* organoids with CRISPR/Cas9 mediated Jak1 knockout (two independent guide RNA’s), Jak2 Knockout (two independent guide RNA’s), Jak1/Jak2 double knockout (Combination of two independent guide RNA’s), a control non targeting guide RNA and Ruxolitinib treated organoids. **G)** Westernblot of JAK-STAT signaling components in LentiCre *Tp53^Δ/Δ^ Rb1^Δ/Δ^* organoids with CRISPR/Cas9 mediated knockout of Jak1 and/or Jak2 (2 separate gRNAs per target). Proteins probed as labeled, Actin was used as a loading control. Proteins as marked.

To interrogate the role of Jak/Stat pathway activation in the initiation of plasticity further, we derived *Trp53^loxP/loxP^, Rb1 ^loxP/loxP^* organoids in which either *Jak1* or *Jak2* alone or in combination was disrupted using CRISPR/Cas9-directed sgRNAs. We then deleted *Trp53* and *Rb1* by lentiviral Cre infection and scored the percentage of organoids with cystic, hyperplastic or slithering phenotypes at 8 weeks, as in **Fig 1B**. Disruption of *Jak1* or *Jak2* alone blunted upregulation of pStat1 and pStat3, but only modestly impacted development of the plasticity phenotype ( a ∼6% to ∼13% (p=0.01, Jak1 guides) and ∼15% (p=0.09 Jak2 guides) increase in cystic organoids and no change in slithering phenotype (∼4% versus ∼6%), with most organoids progressing to the hyperplastic morphology (**Fig 3F-G**). In contrast, organoids with combined *Jak1/Jak2* disruption (which more effectively blocked Stat1/3 phosphorylation) had substantial preservation of cystic morphology (>3-fold increase). This genetic dependence on Jak1 and Jak2 for the development of plasticity raised the question whether pharmacologic intervention might have similar effects, including potential reversal of the organoid morphology that had already progressed to a fully plastic state phenotype.

Indeed, treatment with the dual Jak1/2 kinase inhibitor Ruxolitinib (Rux) restored cystic morphology (from ∼5% to ∼40%) in a dose-dependent fashion (**Fig 3F, S7D**). As further evidence of reversion to a more luminal lineage, these morphological changes were accompanied by a substantial increase in Ar expression, as well as a return of elevated pStat1 and pStat3 levels back to baseline (**Fig 3E**).

Because our prostate organoids are cultured using defined, serum-free conditions that, notably, do not contain IFN or other known ligands that might activate IFN response genes, we explored potential mechanisms as to how the pathway might be stimulated following *Trp53/Rb1* co-deletion. The enrichment of JAK-STAT3 and LIF signaling signatures suggested that IFN response genes may be activated by cytokine receptors, perhaps through an autocrine loop. We reasoned that if a ligand-receptor pair activates JAK-STAT, then high abundance of the ligand-receptor should correlate with increased JAK-STAT signaling in the mutant organoid cells. A key advantage in our organoid model (versus an *in vivo* setting of plasticity), is that we only needed to consider autocrine ligand-receptor (L-R) interactions and were able to exclude cell type as a variable in L-R analysis. We evaluated a set of 74 L-R pairs known to activate JAK-STAT signaling and scored each candidate based on their correlation with a summary statistic of 33 JAK-STAT associated genes enriched following mutation (**Table S12-13;** see **Methods**).

The top ligand-receptor pairs identified by this analysis were LIF and its heterodimeric receptor (LIFR/IL6ST), FGF1/FGFR, IL15/IL15RA, the chemokines CCL2 and CCL5 and their cognate receptors CCR2 and CCR5, and HGF/MET (**Fig 4A, S8A**). Additionally, we reasoned that the ligand-receptor pairs responsible for an increase in plasticity should display: (1) upregulation of receptor expression early in the time course (week 2), and (2) further upregulation with antiandrogen treatment. Based on these criteria, FGFR and LIF (co)receptors remained as top candidate drivers, based on early expression of FGFR1-3 and LIFR following *Trp53/Rb1* co-deletion as well as increased expression of FGFR1, LIFR and IL6ST following ENZ treatment (**Fig 4B, S8C-D**).

**Figure 4:**
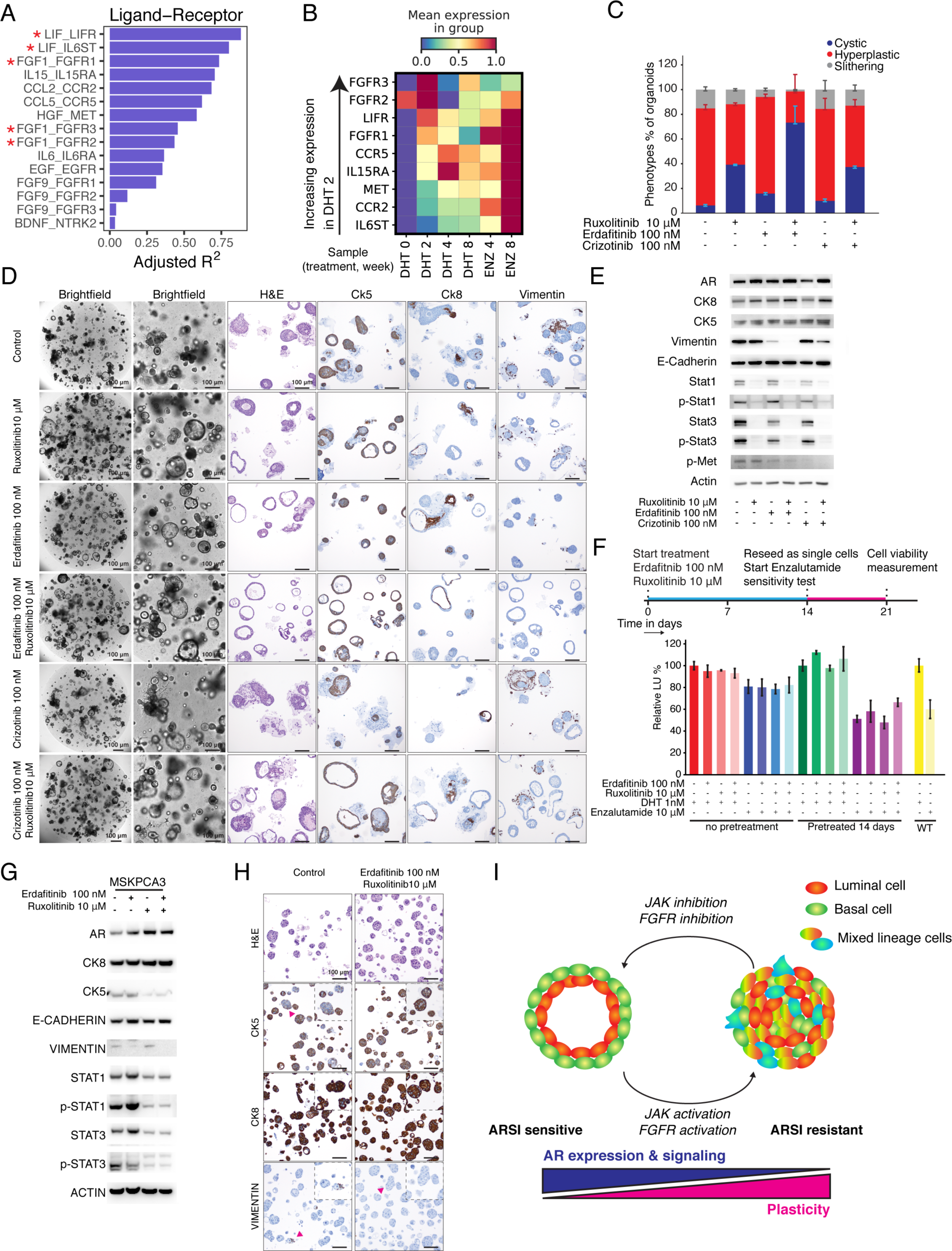
Pharmacological inhibition of JAK-STAT signaling and FGFR signaling resensitizes prostate cancer organoids to ARSI. **A)** Barplot showing the top scoring ligand-receptor (L-R) interactions known to activate JAK/STAT signaling, based on adjusted R^2^ (Radj^2^) of the regression *JAK_STAT ∼ L + R + L:R*, where *JAK_STAT* is the JAK/STAT signaling score, and *L* and *R* represent scaled imputed ligand and receptor expression (see **Methods**). Only L-R pairs with non-zero Radj^2^ are shown. Red asterisks indicate L-R pairs where *R* is overexpressed early at timepoint week 2 (see **Figure S8B**). **B)** Heatmap of mean gene expression (rows) over sample timepoints (columns) for candidate receptors activating downstream JAK/STAT signaling in SEACells. Receptor genes are ordered by increasing expression in the early timepoint at week 2 from bottom to top. **C)** Quantification of phenotypes cystic (Blue), Hyperplastic (Red) and slithering (Silver) of *Tp53^Δ/Δ^ Rb1^Δ/Δ^* organoids treated for 14 days with indicated drugs, see methods for exact medium composition. **D)** Representative brightfield pictures, H&E staining and IHC of CK8, Ck5 and Vimentin of *Tp53^Δ/Δ^ Rb1^Δ/Δ^* organoids treated for 14 days with indicated drugs. Scale bars represent 100 μm. **E)** Western blot of lineage markers and JAK-STAT signaling components in *Tp53^Δ/Δ^ Rb1^Δ/Δ^* organoids treated for 14 days with indicated drugs, see methods for exact medium composition. Proteins probed as indicated. Actin was used as a loading control. **F)** Top: Schematic overview of resensitization drug experiments. Organoids are treated with Erdafitinib 100 nM and Ruxolitinib 10 μM for 14 days in low Egf organoid medium (ENR+A83+DHT), control organoids were cultured in low EGF organoid medium for 14 days. Subsequently organoids (10000 cells per well, triplicate) are reseeded in organoid culture medium without EGF (NR+A83), containing an AR agonist (DHT 1 nM) or antagonist (Enzalutamide 10 μM). Viability was measured by CellTiterGlo after 7 days of Enzalutamide treatment of *Tp53^Δ/Δ^ Rb1^Δ/Δ^* organoids treated for 14 days with indicated drugs (Lower). See methods for exact medium composition. **G)** Westernblot of lineage markers and JAK-STAT signaling components in MSKPCA3 organoid after 14 days treatment with Erdafitinib 100 nM, Ruxolitinib 10 μM or with Erdafitinib 100 nM & Ruxolitinib 10 μM combination. Proteins probed as indicated. Actin was used as a loading control. **H)** Representative histology and IHC (CK8, CK5 and Vimentin) of MSKPCA3 organoids after 14 days treatment with Erdafitinib (100 nM) & Ruxolitinib (10 μM) combination in full organoid medium (ENRFFPN, A83-01, Nicotinamide, DHT). See methods for exact medium composition. Scale bars represent 100 μm. **I)** Proposed model system of lineage plasticity. JAK-STAT signaling activation and FGFR signaling activation leads to a AR^low^, ARSI insensitive state that can be reprogrammed back to an ARSI sensitive state. Potentially the JAK-STAT^high^, AR^low^, ARSI insensitive state is a cellular state preceding AR negative ARSI insensitive CRPC.

As a first test of these two pathways, we added recombinant FGF1 or LIF to organoids following *Trp53/Rb1* co-deletion, reasoning that plasticity might be further enhanced by an artificial increase in ligand concentration. As a control for our prioritization schema, we included recombinant HGF since HGF was among the top candidates in our L-R analysis, but MET was not substantially upregulated early at week 2 following *Trp53/Rb1* co-deletion or after Enz (**Fig 4A-B, S8B-C**). FGF1, but not LIF or HGF, dramatically enhanced the magnitude of slithering (from 6% to 50% of matrigel surface) within 3 days (**Fig S9A**), providing further support for Fgfr signaling as a driver of plasticity.

The availability of a clinical grade FGFR inhibitor (erdafitinib), together with the fact that FGFR activation has been implicated in prostate cancer progression, including AR-negative, SYP-negative disease (*4, 22*), led us to examine the functional consequences of pharmacological FGFR inhibition on plasticity, alone and in combination with JAK inhibition. Although treatment with the FGFR inhibitor erdafitinib (Erda) modestly reduced the percentage of organoids with the slithering phenotype, we observed a striking ∼12-fold increase (5% to 60%) in cystic morphology when fully plastic organoids were co-treated with Erda + Rux (**Fig 4C-D**). Consistent with this dramatic morphologic change, western blot analysis showed near complete loss of expression of the mesenchymal lineage marker Vim and increased levels of the luminal marker Ck8 and Ar (**Fig 4E**). In contrast, the MET inhibitor Crizotinib had minimal activity and did not enhance the anti-plasticity effect of Rux. The remarkable restoration of cystic luminal organoids following combined Fgfr + Jak kinase inhibition led us to ask if sensitivity to antiandrogen therapy is also restored. First, we noted that neither Erda nor Rux, alone or in combination, had significant antiproliferative effects in *Trp53/Rb1*-deleted organoids, despite their substantial effects in restoring cystic morphology. However, once this cystic morphology was restored (after 14 days), Enz sensitivity was restored to previously Enz-resistant organoids (∼50% decrease in proliferation, comparable to wild type organoids) (**Fig 4F, S9B**).

To ask if the *Jak*- and *Fgfr*-dependent plasticity observed in our murine organoid model extends to human prostate cancer, we selected three Enz-resistant patient-derived organoids with combined *TP53/Rb1* deletion to replicate the genetic context of our organoid model (*28*). Combination Rux + Erda treatment of MSKPCA3 organoids, but not MSKPCA1 and MSKPCA6 organoids, led to enhanced expression of AR and CK8 as well as reduced CK5, consistent with reprogramming toward a luminal lineage phenotype, as well as induction of modest sensitivity to Enz (∼20% growth reduction) (**Fig 4G-H, S9C-E**). One potential explanation for the differential sensitivity of the human organoids to Rux + Erda despite shared mutant- *TP53/RB1* genotypes is chromatin accessibility, as suggested by a recent analysis of CRPC tumors (including 2 of the 3 human organoids tested here) that defined 4 distinct subclasses based on transcription factor/transcriptome profiles (*29*). Of note, MSKPCA3 falls into a subclass defined by inflammation, IFN response and EMT pathway signaling, reminiscent of our mouse organoid model, whereas MSKPCA1 maps to a distinct subgroup. In addition, MSKPCA3 and our *Trp53/Rb1*-deleted mouse organoids retain modest levels of AR expression, whereas MSKPCA1 and MSKPCA6 are AR-negative, raising the possibility that reversal of plasticity is only possible in cells with some residual luminal gene expression.

In summary, co-deletion of the Trp53 and Rb1 tumor suppressor genes in normal mouse prostate epithelial cells initiates a program of transcriptional changes that, over a period of weeks, results in mixing of luminal and basal identities and eventual progression to mesenchymal and neuroendocrine lineage phenotypes. We find that execution of this lineage plasticity program is dependent on JAK and FGFR kinase signaling and is reversible through combination therapy with the clinically approved pharmacologic inhibitors Rux and Erda (**Fig 4I**). In the reduced complexity setting of our organoid model (with only epithelial cells present), we identify the IL-6 family cytokine LIF and FGF as the sources of JAK/STAT activation but recognize that microenvironmental stimuli (immune cells) are likely to play a role *in vivo*. Of note, IL-6 has been previously implicated in Ar-negative, pStat3+ tumor progression using cell lines derived from the TRAMP prostate cancer model, where Trp53 and Rb1 function are presumably disabled by expression of SV40 large T antigen (*30*), and LIF expression by pancreatic stellate cells can act as a paracrine factor to promote pancreas cancer progression (*31*). The dependence of tumor cells on JAK/STAT and FGFR signaling to initiate a plasticity program, while compelling from a translational perspective, raises questions about how these pathways intersect with the master regulator transcription factors associated with execution of specific lineage transitions (see accompanying manuscript). Our model suggests that EMT transcription factors are upregulated commensurate with Jak/Stat and Fgfr activation whereas upregulation of canonical neuroendocrine factors (Ascl1, Neurod1) may occur later and/or require an appropriate microenvironmental niche. Whatever the mechanism, our current work has near term clinical relevance because reversion to luminal lineage identity is accompanied by restored sensitivity to ARSIs. Drugs such as Rux and Erda, as well as EZH2 inhibitors (*12, 23*), are all compelling candidates to prevent or overcome ARSI resistance in patients at risk for lineage plasticity assuming, as suggested by our human organoid data, we can develop biomarker tools to identify those tumors most amenable to lineage reversion.

## Acknowledgements

The authors thank the Pe’er lab and the Sawyers lab for valuable critiques and discussions. Molecular Cytology Core Facility from MSKCC for help with confocal microscopy and IHC. Flow Cytometry Core Facility from MSKCC for help with FACS experiments.

## Data accession

All sequencing data is deposited at the Gene Expression Omnibus (GEO) database.

## Author contributions

W.R.K, J.C, D.P and C.L.S. conceived the project. W.R.K and C.L.S. designed organoid experiments. W.R.K, J.R.L and S.Z. performed organoid experiments and downstream benchwork. J.C and D.P. designed computational approach. J.C., M.S and D.P. developed computational methods. Z.C., S.P. and D.P. contributed computational methods. J.C. and M.S. performed computational analysis. J.R.L., J.L., K.E.L. and W.R.K. performed mouse work. I.M., O.C. and X.T performed single cell experiments. L.M. and R.C oversaw single cell experiments. D.P., C.L.S, J.C, W.R.K wrote the manuscript. D.P. and C.L.S. oversaw the project.

## List of supplementary materials

**Supplementary Figure 1.**
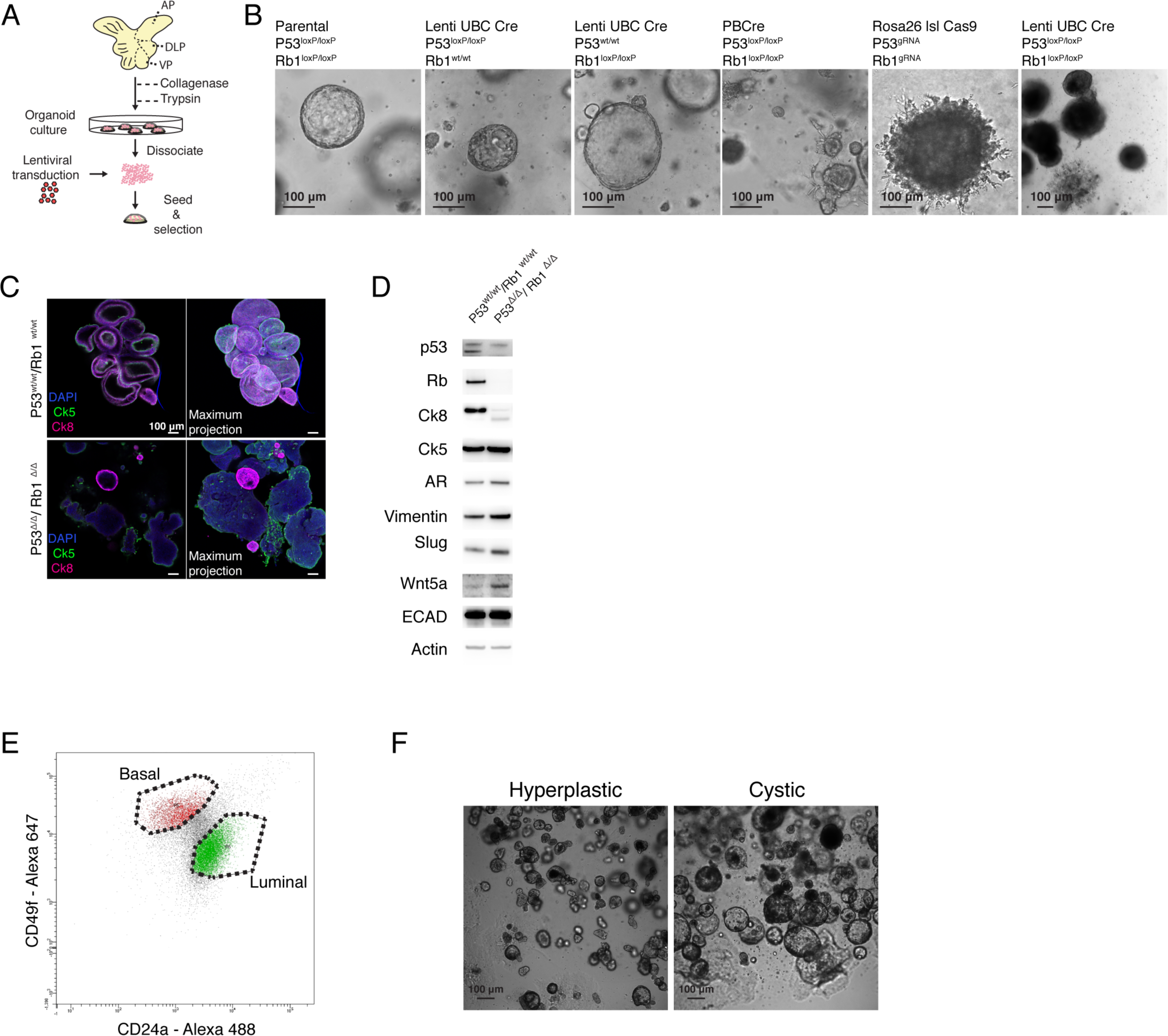
Establishment of an organoid model of spontaneous lineage plasticity. **A)** Schematic overview of organoid establishment followed by lentiviral mediated recombination of *p53* and *Rb1* loci. **B)** Representative brightfield pictures of wild-type, single knockout *Tp53*, Single knockout *Rb1*, PBCre *Tp53^Δ/Δ^ Rb1^Δ/Δ^* organoids, CRISPR/Cas9 mediated *Tp53^Δ/Δ^ Rb1^Δ/Δ^* organoids and LentiCre *Tp53^Δ/Δ^ Rb1^Δ/Δ^* organoids. All organoids were cultured for > 3 months. **C)** Representative immunofluorescent staining of CK5 (green) and CK8 (Magenta) *Tp53^loxP/loxP^ Rb1^loxP/loxP^* organoids) and ∼10 weeks post deletion (LentiCre *Tp53^Δ/Δ^ Rb1^Δ/Δ^* organoids), single Z-scan and maximum projection. Scale bars represent 100 μm. **D)** Representative western blot of wild-type (*Tp53^loxP/loxP^ Rb1^loxP/loxP^* organoids) and ∼10 weeks post deletion (LentiCre *Tp53^Δ/Δ^ Rb1^Δ/Δ^* organoids). Proteins as marked **E)** Representative FACS plot and gating of CD24 and CD49f expressing cells in organoids. **F)** Representative picture of hyperplastic and cystic cultures 4 weeks post CRISPR/Cas9 mediated *Tp53* and *Rb1* deletion/mutation. Scale bars represent 100 μm.

**Supplementary Figure 2.**
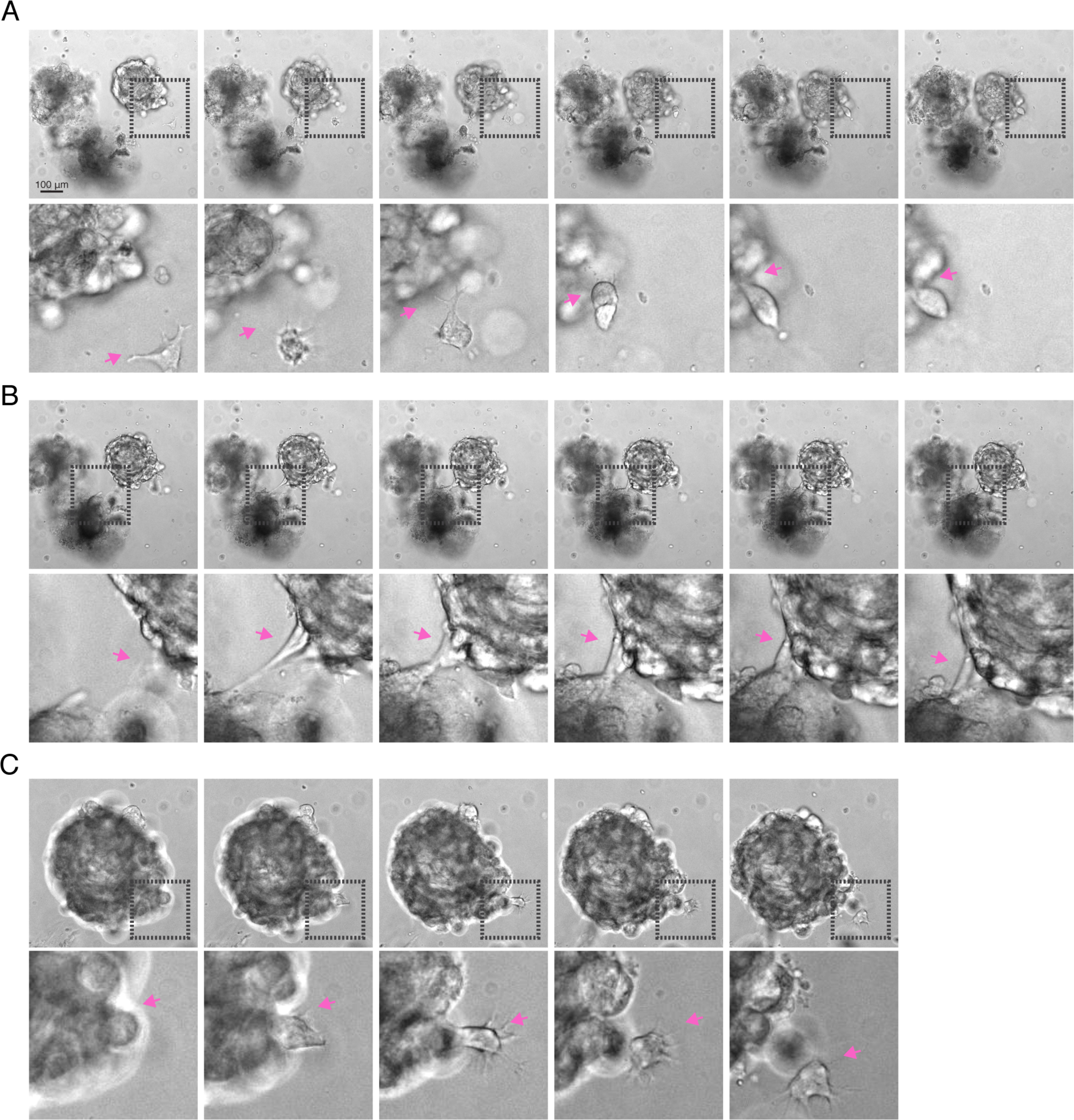
Establishment of an organoid model of spontaneous lineage plasticity. **A, B, C)** Stills of 7-day movies of LentiCre *Tp53^Δ/Δ^ Rb1^Δ/Δ^* organoids (∼10 weeks post Cre recombination) showing high motility and plastic behavior.

**Supplementary Figure 3.**
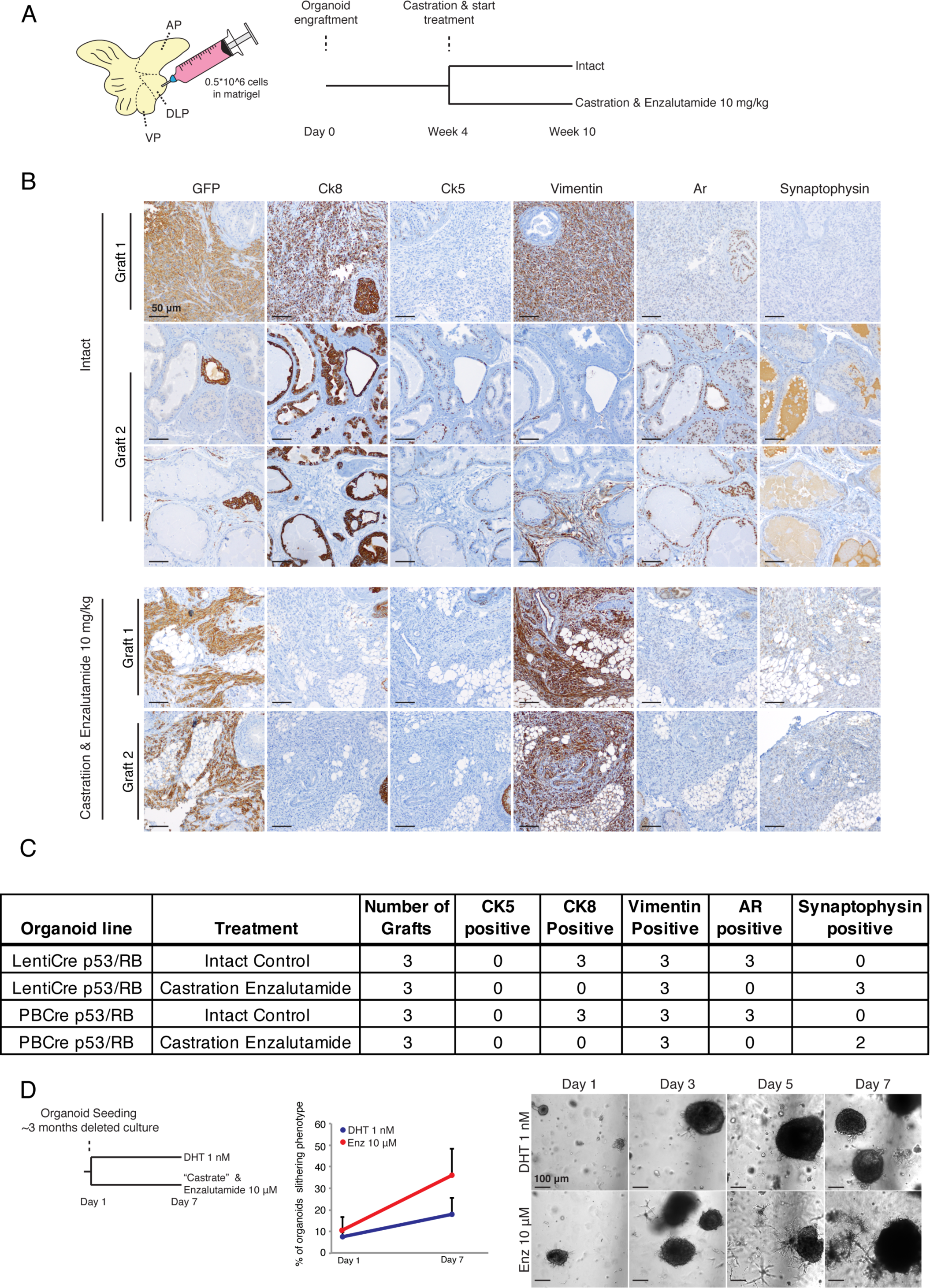
Establishment of an organoid model of spontaneous lineage plasticity: **A)** Schematic overview of orthotopic transplantation of 5*10~5 cells derived from *Tp53^Δ/Δ^ Rb1^Δ/Δ^* organoids (∼10 weeks post Cre recombination). Castration and treatment with Enzalutamide (10 mg/kg) was started 4 weeks post organoid engraftment. **B)** Representative IHC of GFP, Ck8, Ck5, Vimentin, Ar and Synaptophysin in orthotopic grafts of LentiCre *Tp53^Δ/Δ^ Rb1^Δ/Δ^* organoids 10 weeks post engraftment, with and without 6-week treatment of castration and Enzalutamide (10 mg/kg). GFP staining was used to identify graft. Scale bars represent 50 μm. **C)** Table showing IHC patterns of Ck8, Ck5, Vimentin, Ar and Synaptophysin in orthotopic grafts of LentiCre *Tp53^Δ/Δ^ Rb1^Δ/Δ^* organoids or PBCre *Tp53^Δ/Δ^ Rb1^Δ/Δ^* organoids in intact hosts and castrated hosts treated with Enzalutamide (10 mg/kg). **D)** Left: Schematic overview of experimental setup of ARSI treatment *in vitro*. Middle: Quantification of slithering phenotype in organoids cultured for 7 days with agonist (DHT, 1nM DHT) and antagonist (Enzalutamide, 10 μM) of AR. Right: Representative brightfield pictures of organoid cultures during treatment. Scale bars represent 100 μm.

**Supplementary Figure S4.**
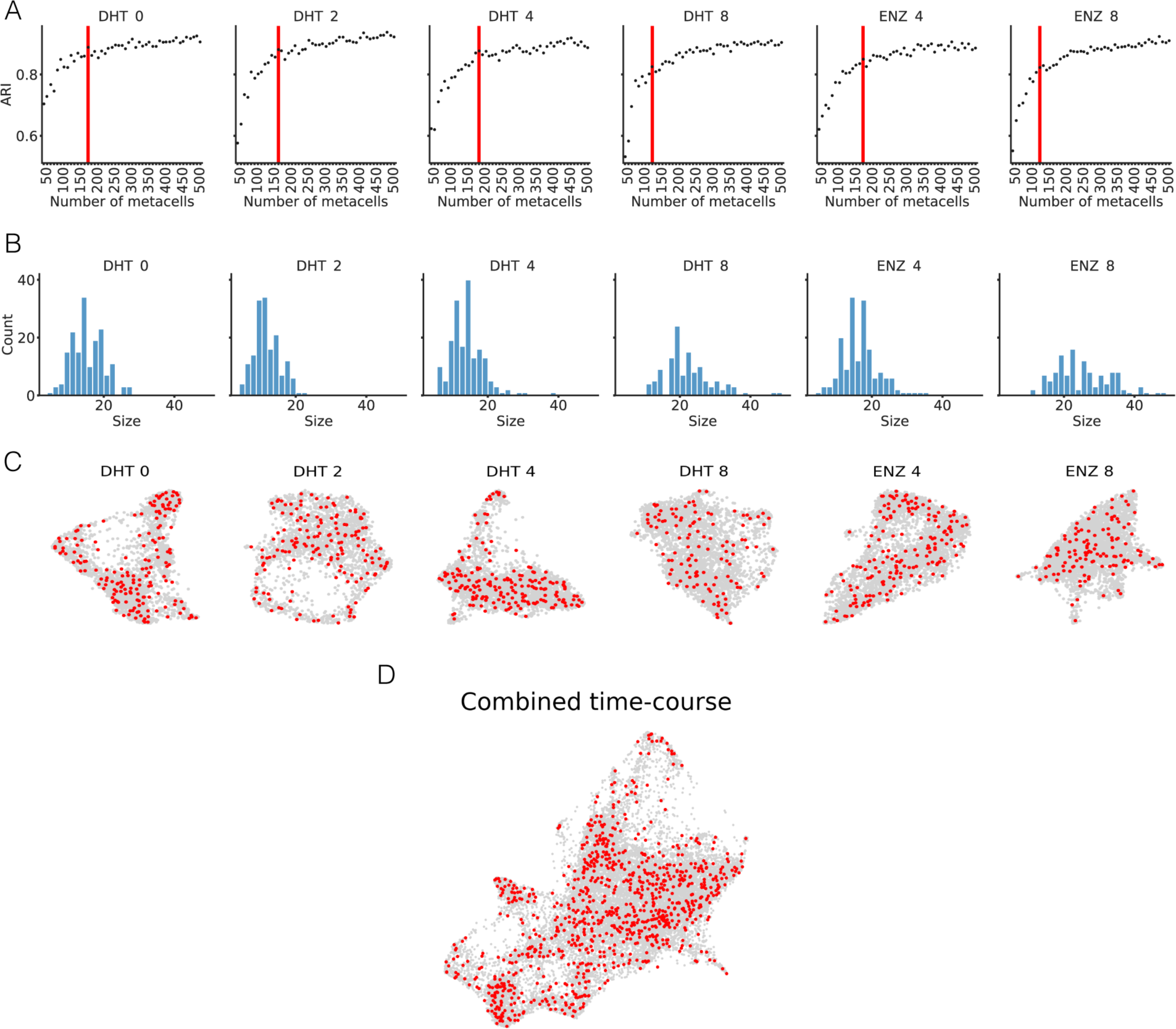
**A)** Robustness of SEACell metacells between consecutive values of *m* archetypes, as assessed by adjusted Rand index ARI. Red line indicates kneepoint of the ARI curve, indicating the minimum *m* archetypes beyond which further increase in robustness plateaus. **B)** Kernel density plots of metacell size distribution. SEACell archetypes (red) projected on force-directed layouts of single cells **(C)** within each sample and **(D)** in the combined dataset. Each red dot corresponds to the single cell closest to the centroid of a given SEACell archetype.

**Supplementary Figure S5.**
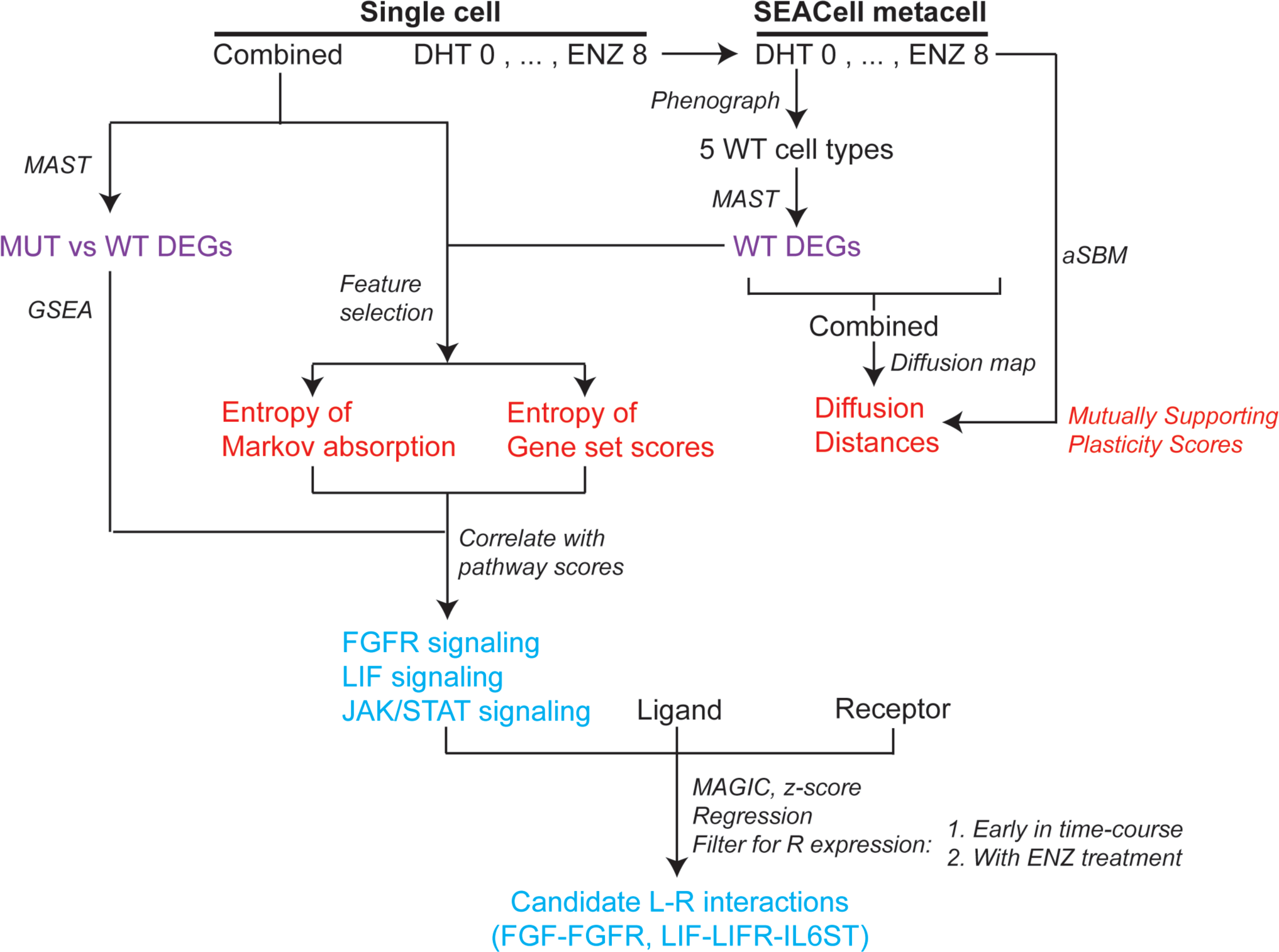
Schematic of the computational analysis of the scRNA-seq dataset from the *Rb1/Trp53*-deleted prostate organoid.

**Supplementary Figure S6.**
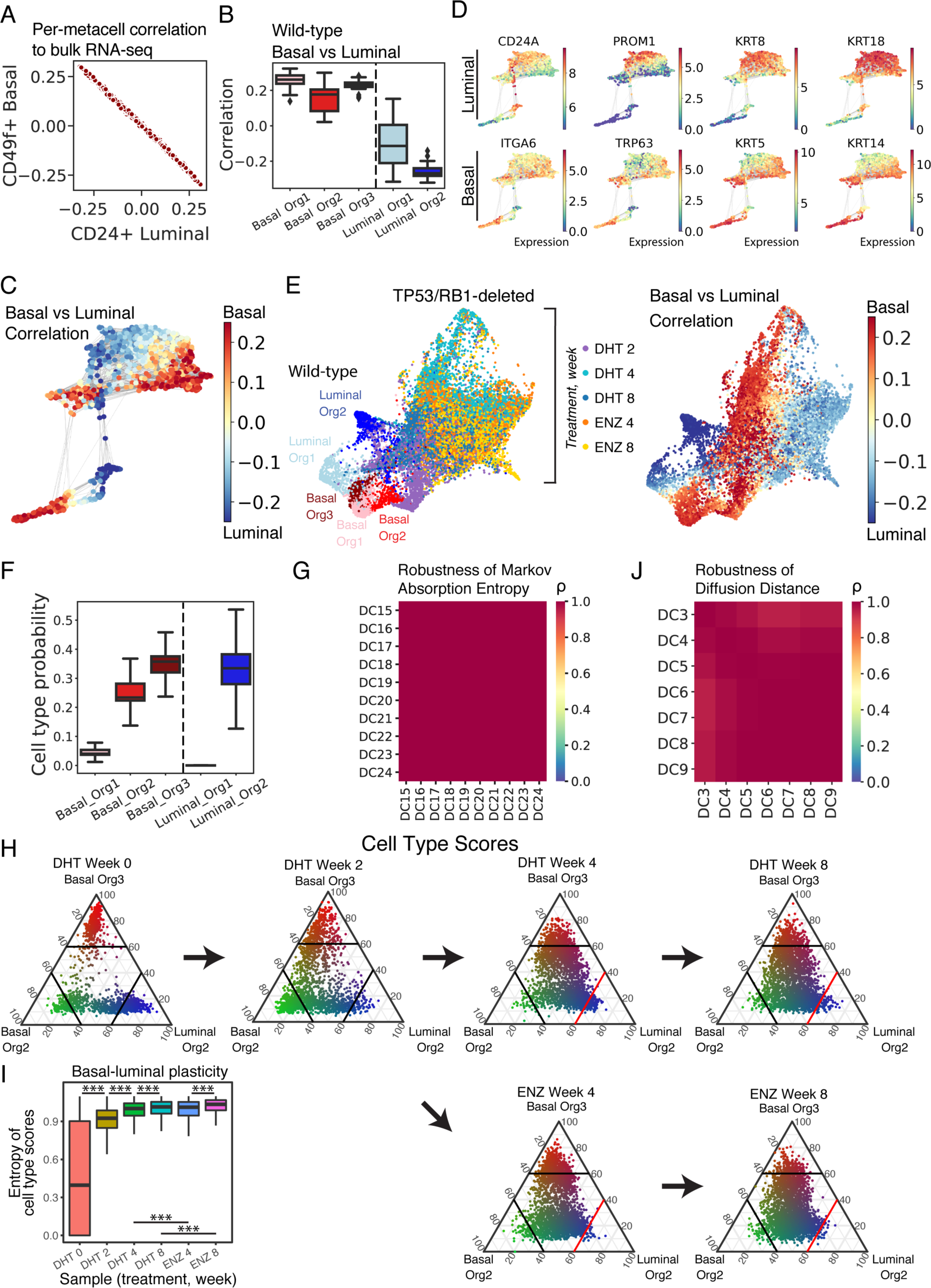
**A)** Scatterplot showing the Pearson’s correlation of z-scored expression per metacell in the combined mutant and wild-type dataset to bulk-sorted CD24+ luminal cells vs CD49f+ basal cells. Genes are restricted to significant DEGs for each wild-type cell type, with gene sets non-specific to cell type removed (i.e. cell cycle, apoptosis, hypoxia; see **Methods**). **B)** Boxplot of correlation to bulk-sorted CD49f^+^ basal vs CD24^+^ luminal metacells for each basal and luminal subpopulation in the wild-type organoid. **C)** Force-directed layout (FDL) of metacells in the prostate organoid cells before and after *Rb1/Tp53* deletion, colored by correlation to bulk-sorted basal CD49f^+^ and luminal CD24^+^ cells (same layout as **Fig 2D**). Edges indicate k-nearest neighbors (k=10). **D)** FDL of metacells in the prostate organoid cells before and after *Rb1/Trp53* deletion, showing gene expression of basal and luminal markers. Expression is normalized, and log2 transformed with pseudocount of 1. Edges indicate k-nearest neighbors (k=10). **E)** FDL of single cells in the prostate organoid cells before and after *Rb1/Tp53* deletion, annotated by (left) wild-type basal and luminal subpopulations as well as time and treatment following mutation; and (right) correlation to bulk-sorted basal CD49f^+^ and luminal CD24^+^ cells. The FDL is constructed using a kNN graph, where k = 15 (see **Methods**). **F)** Boxplot of cell type probabilities for single cells in the mutant organoid. Cell type classification probabilities are calculated by Markov absorption in diffusion maps using wild-type basal and luminal subpopulations as training labels (see **Methods**). **G)** Robustness of the entropy of classification probabilities for each cell type as a measure of plasticity, using Pearson’s correlation (ρ). This *per-*cell plasticity score does not vary substantially as the number of DCs changes from 15 to 24. **H)** Ternary plots of cell-type probabilities based on gene set scores for single cells in the organoid before and after mutation. Each ternary plot represents a timepoint/condition. Cell-type probabilities are calculated as the average Z-score of DEGs for each of the predominant basal and luminal organoid subtypes in the mutant setting (Basal Org2, Basal Org3, and Luminal Org2) (see **Methods**). Each dot is a cell plotted using 3 coordinates representing the probabilities of representing the 3 most represented basal and luminal phenotypes in the wild-type organoid: Basal Org2 (green), Basal Org3 (red), and Luminal Org2 (blue). Color intensity of each dot corresponds to the certainty of representing a given cell type, calculated by entropy. Higher entropy indicates greater uncertainty of cell type, increased basal-luminal mixing, and increased plasticity. The red line at 60% probability of Luminal Org2 classification at weeks 4 and 8 highlights the decreased probability for Luminal Org2 in Enz-treated samples compared to DHT control **I)** Boxplot of entropy of cell-type probabilities for single cells based on gene set scores, as a measure of plasticity over time and treatment (Wilcoxon rank-sum test, see **Methods**). p-values: ***<0.001. **J)** Robustness of log mean diffusion distance between basal and luminal metacells as a measure of plasticity, using Pearson’s correlation (ρ). This *per-*cell plasticity score does not vary substantially as the number of DCs changes from 3 to 10.

**Supplementary Figure S7.**
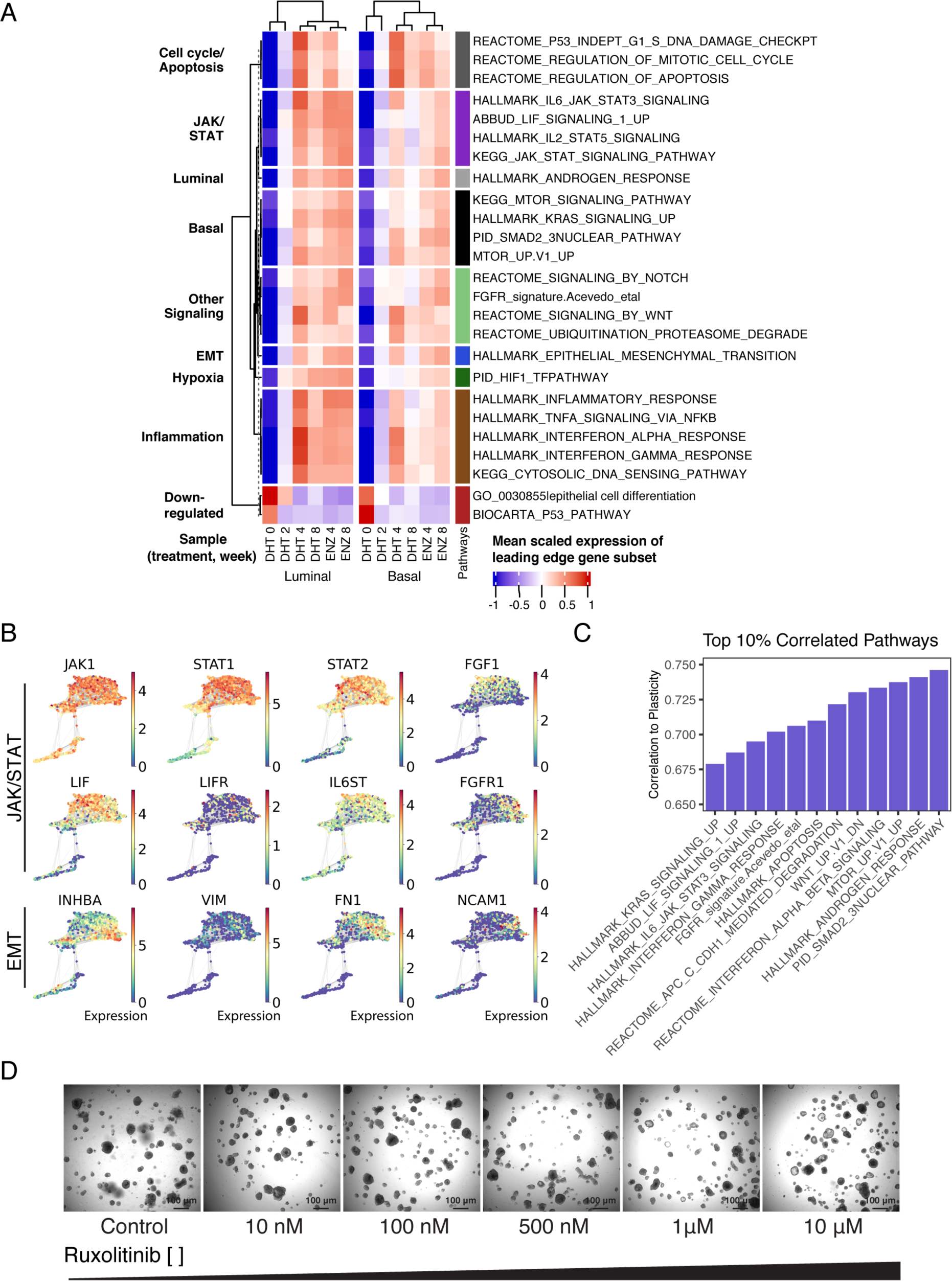
**A)** Heatmap showing enriched pathways in each timepoint stratified by basal and luminal cell types. Each pathway score is measured as the average Z-score of gene expression in the leading-edge gene set of each pathway among SEACells (see **Methods**). **B)** FDL of SEACells in the prostate organoid cells before and after *Rb1/Trp53* deletion (same layout as **Fig 2D**), showing gene expression of important markers for JAK/STAT signaling and EMT. Expression is normalized, and log2 transformed with pseudocount of 1. **C)** Barplot showing select pathways from the top 10% correlated to plasticity. The full list of the top 10% pathways is shown in **Table S11**. Plasticity is measured by entropy of cell-type classification probabilities in single cells. Each pathway is scored by the mean expression of the leading-edge subset of each pathway. Pearson’s correlation is then calculated for each pathway score and plasticity (see **Methods**). **D)** Representative brightfield pictures of organoids treated with increasing doses of the Janus Kinase inhibitor Ruxolitinib (0, 10 nM, 100 nM, 10 μM, 10 μM) 1 for 21 days. **H)** Representative histology and IHC (CK8, CK5 and Vimentin) of MSKPCA3 organoids after 14 days treatment with Erdafitinib (100 nM) & Ruxolitinib (10 μM) combination in full organoid medium (ENRFFPN, A83-01, Nicotinamide, DHT). See methods for exact medium composition. Scale bars represent 100 μm. **I)** Proposed model system of lineage plasticity. JAK-STAT signaling activation and FGFR signaling activation leads to a AR^low^, ARSI insensitive state that can be reprogrammed back to an ARSI sensitive state. Potentially the JAK-STAT^high^, AR^low^, ARSI insensitive state is a cellular state preceding AR negative ARSI insensitive CRPC.

**Supplementary Figure 8:**
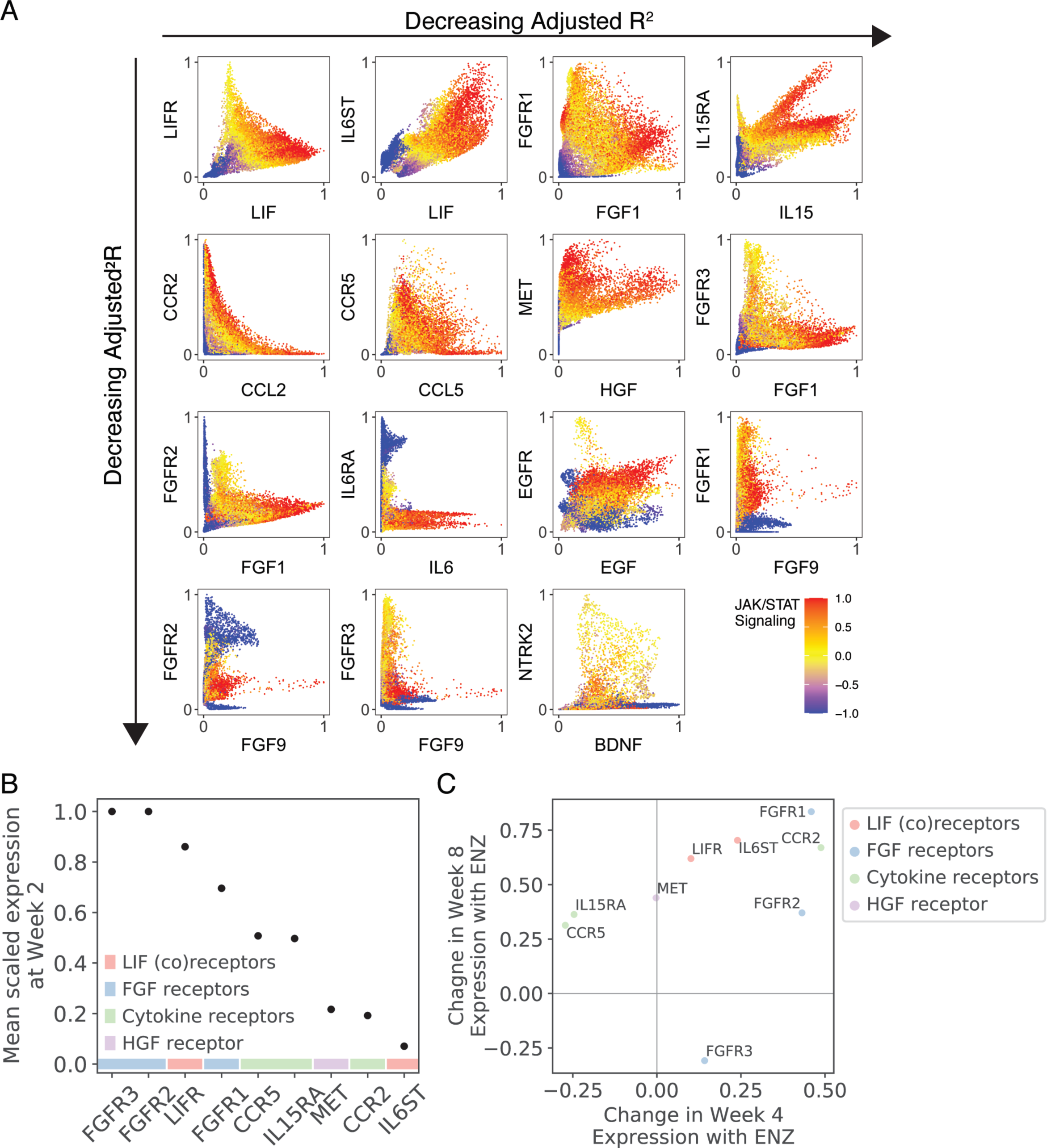
**A)** Scatterplots of top ligand-receptor interactions known to activate JAK/STAT signaling. Each dot represents a single cell plotted by Z-scored, imputed ligand vs receptor expression, colored by JAK/STAT signaling score. Scatterplots are ordered by decreasing adjusted R^2^ from left to right, then top to bottom (**see Methods**). **B)** Scatterplot of the mean scaled gene expression at early timepoint week 2 for candidate receptors leading to downstream JAK/STAT signaling among SEACells, with LIF (co)receptors and FGF receptors having the highest expression. Mean expression is scaled across all sample timepoints from 0 to 1. Receptor genes are annotated by color based on the corresponding class of ligands. **C)** Scatterplot of change in expression with ENZ treatment at week 4 (x-axis) and week 8 (y-axis) for candidate receptors leading to downstream JAK/STAT signaling among SEACells. Receptor genes are annotated by color based on the corresponding class of ligands. LIF (co)receptors, FGF receptors, and CCR2 show increased expression with ENZ treatment at both week 4 and 8.

**Supplementary Figure 9:**
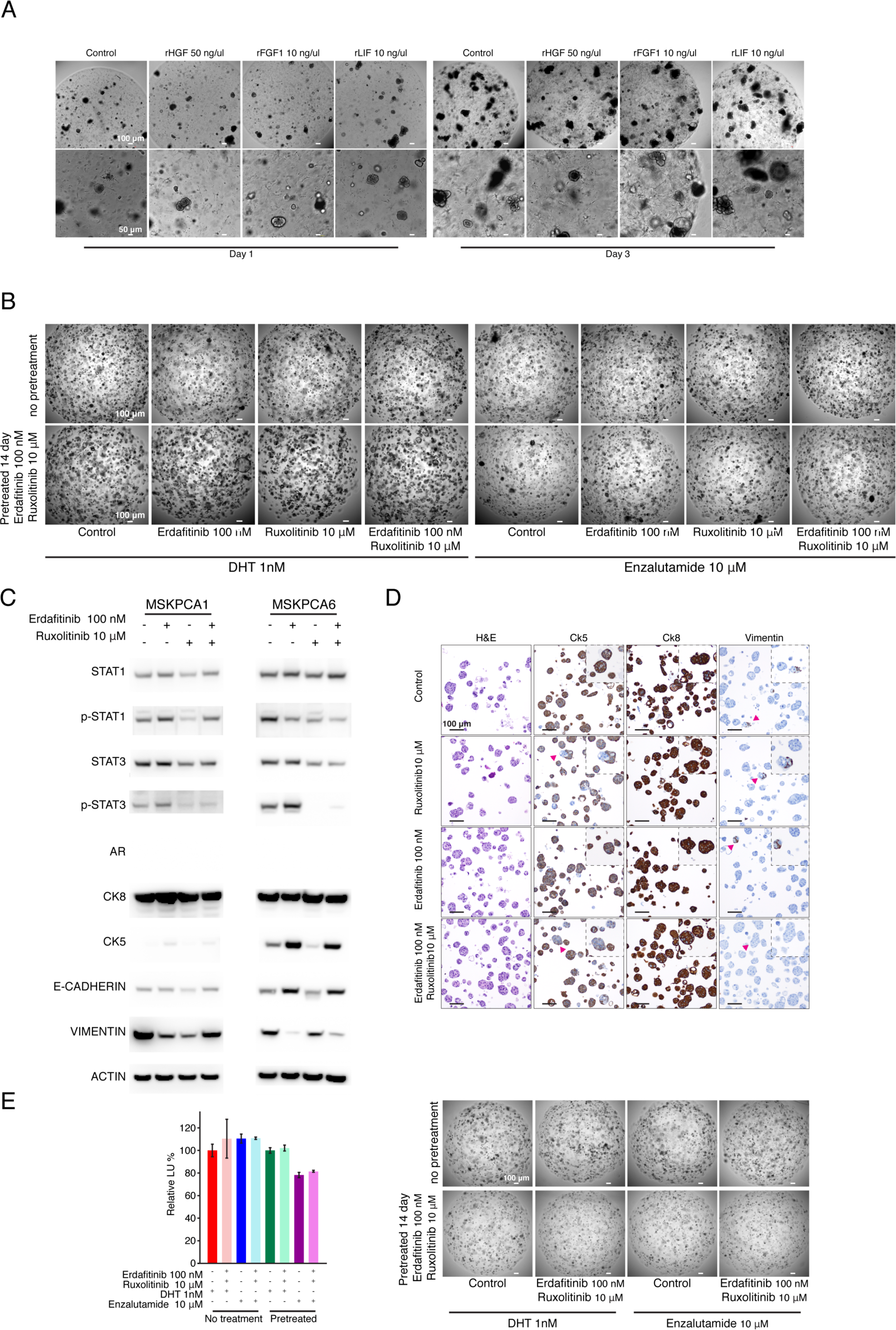
Pharmacological inhibition of JAK-STAT signaling and FGFR signaling resensitizes prostate cancer organoids to ARSI. Related to figure 4. **A)** Representative brightfield pictures of *Tp53^Δ/Δ^ Rb1^Δ/Δ^* organoids treated for 7 days with recombinant HGF (50 ng/ml), FGF1 (10ng/ml) and LIF (10 ng/ml). **B)** Representative brightfield pictures of *Tp53^Δ/Δ^ Rb1^Δ/Δ^* organoids after 7 days of DHT (1 nM) Enzalutamide (10 μM) treatment post 14 days treatment with Erdafitinib 100 nM, Ruxolitinib 10 μM or with Erdafitinib 100 nM & Ruxolitinib 10 μM combination. **C)** Westernblot of MSKPCA1 and MSKPCA6, two organoid lines with a double negative CRPC phenotype i.e., AR and Neuroendocrine marker negative, after 14 days treatment with Erdafitinib 100 nM, Ruxolitinib 10 μM or with Erdafitinib 100 nM & Ruxolitinib 10 μM combination. Proteins probed as indicated. **D)** Representative histology of MSKPCA3 organoids after 14 days treatment with Erdafitinib (100 nM), Ruxolitinib (10 μM) or with Erdafitinib (100 nM) & Ruxolitinib (10 μM) combination in full organoid medium (ENRFFPN, A83-01, Nicotinamide, DHT). Images of control and Ruxolitinib & Erdafitinib organoids are the same as in **Figure 4H**. **E)** Viability as measured by cell titer glo. MSKPCA3 Organoids are treated with Erdafitinib 100 nM and Ruxolitinib 10 μM for 14 days in organoid medium (ENRFFPN, A83-01, Nicotinamide, DHT). Subsequently organoids (10000 cells per well, triplicate) are reseeded in organoid culture medium without EGF (PNR+A83-01, Nicotinamide), containing an AR agonist (DHT 1 nM) or antagonist (Enzalutamide 10 μM). Viability was measured by CellTiterGlo after 7 days of Enzalutamide treatment of PCa organoids treated for 14 days with indicated drugs (Lower). Representative brightfield pictures of MSKPCA3 organoids after 7 days of DHT (1 nM) Enzalutamide (10 μM) treatment post 14 days treatment with or without Erdafitinib 100 nM & Ruxolitinib 10 μM. See methods for exact medium composition.

Table S1 BULK RNA sequencing table

Table S2 QC statistics single cell RNA-seq

Table S3 Differentially expressed genes wild-type organoid cluster_0

Table S4 Differentially expressed genes wild-type organoid cluster_1

Table S5 Differentially expressed genes wild-type organoid cluster_2

Table S6 Differentially expressed genes wild-type organoid cluster_3

Table S7 Differentially expressed genes wild-type organoid cluster_4

Table S8 Differentially expressed genes wild-type

Table S9 Differentially expressed genes wild-type versus mutant

Table S10 Pathway enrichment wild-type versus mutant

Table S11 Pathway activation correlation to plasticity

Table S12 Curated list of JAK-STAT activating Ligand-Receptor pairs

Table S13 JAK-STAT signature

Table S14 Organoid culture conditions

Table S15 guide RNA’s used for CRIPSR/Cas9 mediated knockout

Table S16 Antibodies GMT file: Gene sets curated from MSigDB and literature, used for pathway enrichment analysis Movie S1 Organoid 7 days post deletion Movie S2 Organoid 10 weeks post deletion ** References 32-56 only used in supplementary material section

### Methods

#### Prostate isolation and organoid culture

Murine prostate organoids were established and cultured as described previously(*1,2*). Briefly, prostates were harvested from adult mice harboring the following genotypes. 1) *Tp53^loxP/loxP^* 2) *Rb1^loxP/loxP^* 3) *Tp53^loxP/loxP^, Rb1^loxP/loxP^* 4) *PBcre Tp53^loxP/loxP^, Rb1^loxP/loxP^* (all genotypes described in detail in Ku et al. 2017(*3*) and 5) Rosa26 lsl Cas9(*4*). Prostates were digested with Collagenase Type II (Gibco) for 2 h at 37 °C, followed by TrypLE (Gibco) digestion at 37 °C until a single cell suspension was obtained. Digestions were supplemented with Y-27632 (10 μM) to inhibit anoikis.

Bulk isolated prostatic epithelial cells were embedded in 35-µl drops of basement membrane extract (Matrigel, Corning) and overlaid with mouse prostate organoid medium. Organoids were passaged on a weekly basis using trituration with a fire polished glass Pasteur pipet or by trypsinization using TRYPLE. Human prostate organoids were cultured as described previously(*1,2,5*). A full list of media ingredients and concentrations is given in **Table S14**.

#### P53 and RB1 deletion

Cre-mediated recombination of *Tp53* and *Rb1* loci was performed by lentiviral or adenoviral delivery of CRE recombinase as described previously(*1,6*). CRISPR/Cas9-mediated gene knockout was performed by lentiviral delivery of plasmids encoding Cas9 and gRNA sequences to organoids as described previously(*1,6*) using LentiGuide or LentiCRISPRv2 plasmids(*7*).

#### Drug treatments for redifferentiation

Organoids were treated for 14 d using media compositions as described in **Table S14**. Organoids were subsequently either isolated for histology, protein analysis or seeded in enzalutamide sensitivity assays.

#### Growth and viability assays

Cell growth and viability assays were performed as described previously(*8,9*). Briefly, 5,000 to 10,000 cells were seeded in triplicate and treated as described in **Table S14**. After 7 d, organoids were imaged, and viability was measured using CellTiterGlo.

#### Phenotypic quantification

Organoid phenotypes were quantified 7 d post mechanical splitting via trituration. Phenotypes were binned in three categories: Cystic, Hyperplastic and Slithering.

#### Cloning

Oligonucleotides were cloned into LentiCRISPRv2 (LCV2) and LentiGuide (LG) vectors as described previously(*7*). A full list of oligo sequences and corresponding gene targets are given in Table S15.

#### Mouse work

All animal work was performed in compliance with the guidelines of the Research Animal Resource Center of Memorial Sloan Kettering Cancer Center (IACUC: 06-07-012). For orthotopic injections, NOD SCID Gamma (NSG) mice were purchased from Jackson laboratories. Approximately 500,000 cells derived from *Tp53^Δ/Δ^ Rb1^Δ/Δ^* organoids 10 weeks post deletion were resuspended in 20 µl of Matrigel: Organoid Medium (1:1) mix. Cells were injected into the dorsal lobe of the prostate in 8-week-old mice. Mice were castrated and treated with Enzalutamide (10 mg/kg) 4 weeks post transplantation as described previously(*10*). Prostates were isolated and processed for downstream analyses.

#### Immunohistochemistry and immunofluorescence

Prostates were fixed with 4% PFA, dehydrated with ethanol and paraffin embedded according to standard protocol. 4-µm slides were cut and placed on glass slides. Immunohistochemistry or immunofluorescence of FFPE tissue was performed using a Ventana BenchMark Ultra. Antibody concentrations and antigen retrieval is listed in **Table S16**. FFPE stained tissue was scanned using a MIRAX scanner and processed using Caseviewer software. For immunofluorescent staining of wholemount organoids, organoids were fixed with 4% PFA on ice for 1 h, then blocked and permeabilized using 1% triton-X100 and 1% FCS in PBS0 for 1 hour at RT. Staining was performed in 0.3% Triton X-100 and 0.0.5% FCS in PBS0 overnight at 4 °C on gently shaking. Stained organoids were imaged using a Leica SP5 or SP8 confocal microscope. Images were processed using Leica software or FIJI software.

#### RNA isolation, cDNA Synthesis and quantitative PCR

RNA was isolated from organoids using the PureLink RNA mini kit (ThermoFisher) according to manufacturer’s protocol. RNA concentration was quantified using a NanoDrop (ThermoFisher).

#### Bulk RNA-sequencing and analysis

RNA sequencing was performed in collaboration with the New York Genome Center. For generation of bulk RNA-seq basal and luminal signatures, basal (CD49f^+^) and luminal (CD24^+^) cells were sorted from three independent prostate samples of *FVB/NJ* mice. For transcriptional profiling of the organoids, three replicates of *Tp53^loxP/loxP^, Rb1^loxP/loxP^* and *Tp53^Δ/Δ^ Rb1^Δ/Δ^* organoids (LentiCre, 10 weeks post deletion) were used. For library preparation, 100 ng (Basal/Luminal) or 1000 ng (Organoids) RNA was prepared using the mRNA TruSeq stranded 175-bp library preparation kit (Illumina). Libraries were sequenced on an illumina HiSeq to a depth of 40 million reads. Raw data was processed and analyzed using Partek Flow software. Differentially expressed genes were generated based on transcripts per million (TPM) used as input to the DESeq2 algorithm using default parameters within the Partek Flow suite.

#### Protein isolation and western blot analysis

Organoids were isolated from the basement membrane extract by multiple washes with ice cold PBS0 in a 15-ml Falcon tube. Cells were lysed in RIPA buffer containing protease inhibitors (Calbiochem) and phosphatase inhibitors (Calbiochem) on ice and sonicated three times for 30 s at 30-s intervals using a Bioruptor. Protein concentrations were quantified using a bicinchoninic acid (BCA) assay (Pierce, Thermo Fisher). Lysates were denatured using 4X protein loading dye (SDS 200 nM Tris, 8% SDS, 0.4% bromophenol blue, 40% glycerol, 400 mM 2-mercaptoethanol, pH 6.8). 5-10 µg of protein was loaded on NuPage 4-12% gradient polyacrylamide gels (Invitrogen). After electrophoresis, protein was transferred to a PVDF membrane and blocked with either 5% milk or 5% BSA in TBS-T. Primary antibodies were incubated overnight. Membranes were washed using TBS-T and incubated with secondary antibodies for 1 h at room temperature. Proteins were visualized using ECL and ECL prime (Amersham, GE healthcare) and ImageQuant 800 (Amersham, GE healthcare). Antibody concentrations and corresponding blocking methods are given in **Table S16**.

#### Single-cell isolation and scRNA-seq

For single-cell analysis, organoids were isolated from basement membrane extract and digested to single-cell suspension by incubating with TRYPLE (Life technologies) supplemented with Y-27632 (10 μM) for 10 min at 37 °C with shaking, followed by a 2-min incubation with EDTA (1 mM), Y-27632 (10 μM) in PBS0 and a second digestion with TRYPLE (Life technologies) supplemented with Y-27632 for 5 min at 37 °C with shaking. Cells were filtered and resuspended in PBS0, Y-27632 (10 μM) and BSA (0.04%) before processing further for single-cell sequencing. Cells were checked for viability using 0.2% (w/v) Trypan Blue staining (Countess II) and all sequencing experiments were performed on samples with a minimum of 80% viable cells.

Single-cell encapsulation and scRNA-seq library prep of FACS-sorted cell suspensions was performed on the Chromium instrument (10x Genomics) following the user manual (Reagent Kit 3’ v2). Each sample loaded onto the cartridge contained approximately 5,000 cells at a final dilution of ∼500 cells/µl. Transcriptomes of encapsulated cells were barcoded during reverse transcription and the resulting cDNA was purified with DynaBeads, followed by amplification per the user manual. Next, PCR-amplified product was fragmented, A-tailed, purified with 1.2X SPRI beads, ligated to sequencing adapters and indexed by PCR. Indexed DNA libraries were double-size purified (0.6–0.8X) with SPRI beads and sequenced on an Illumina HiSeq (R1 – 26 cycles, i7 – 8 cycles, R2 – 70 cycles or higher) to a depth of >50 million reads per sample (>13,000 reads/cell).

#### Pre-processing of scRNA-seq data

FASTQ files from sequenced samples were individually processed using the SEQC pipeline(*11*) based on the mm38 mouse genome reference and default parameters for the 10x single-cell 3’ library. The SEQC pipeline performs read alignment, multi-mapping read resolution, as well as cell barcode and UMI correction to generate a (cell x gene) count matrix. The pipeline performs the following initial cell filtering steps: true cells are distinguished from empty droplets based on the cumulative distribution of total molecule counts; cells with a high fraction of mitochondrial molecules are filtered (>20%), as are cells with low library complexity (cells that express very few unique genes). We used the EmptyDrops package(*12*) to perform additional filtering of empty droplets by retaining cells with FDR < 0.01. Putative doublets were removed using the DoubletDetection package (DOI 10.5281/zenodo.2658729). Genes that were expressed in more than 10 cells were retained for further analysis. The combined dataset yielded a filtered count matrix ! of 15,677 cells by 14,617 genes, with a median of 4,336 transcripts per cell and a median of 2,660 cells per sample.

#### Analysis of single-cell transcriptomes

We used sctransform(*13*) to normalize gene expression in cells pooled across sample timepoints. Given the set of cells *C* and genes *Z*, this method considers the UMI count matrix *U* ∈ ℤ*^C^*^×^*^Z^* to perform a regularized negative binomial regression that calculates a matrix of Pearson residuals *R* ∈ ℝ*^C^*^×^*^Z^*, adjusted for technical artifacts from cellular sequencing depth. This source of bias can be significant in datasets that have variable library size driven by increased proliferation across wildtype and mutant conditions. We have found sctransform normalization to be more faithful than simple library size normalization, particularly in cases where the data consists of similar cell types (e.g. organoid data). For downstream analysis, we use *R* for computations related to the structure of the data, such as principal components analysis (PCA), clustering, and subsequent layouts for visualization and diffusion map construction.

PCA was performed on & (after removing mitochondrial and ribosomal genes) with the top 80 PCs retained with 38% variance explained. To visualize cell layouts, we construct an affinity graph *A* that follows the procedure used in MAGIC(*14*). We first consider the k-nearest neighbors (kNN) graph based on Euclidean distance in PCA space (kNN = 30). We then create affinity graph *A* by applying an adaptive Gaussian kernel (kernel width σ = 10) to the kNN graph that accounts for differential densities of cells in the phenotypic manifold, followed by symmetrizing the matrix. Based on *A*, we calculate the force-directed layout (FDL) using the draw graph function in scanpy(*15*).

To visualize gene expression projected on cell layouts, we normalized the UMI counts ! by library size, scaled by median library size, and log2-transformed with a pseudocount of 1. We chose to perform log transformation to be consistent across our other scRNA-seq study in normal mouse prostate(*9*) and our companion paper investigating plasticity in a transgenic mouse model with *Rb1/Trp53/Pten* deletion, facilitating mapping and reproducibility across datasets. Given the sparse nature of single-cell sequencing that arises from gene dropout, we applied gene imputation with MAGIC (kNN = 30, t = 3)(*14*).

In addition to the combined dataset, we analyzed single cells within each sample timepoint. For each sample *s*, we subset the Pearson residuals to the cells in *s* to *R_s_* and remove mitochondrial and ribosomal genes before performing PCA, obtaining the following PC loadings per sample:

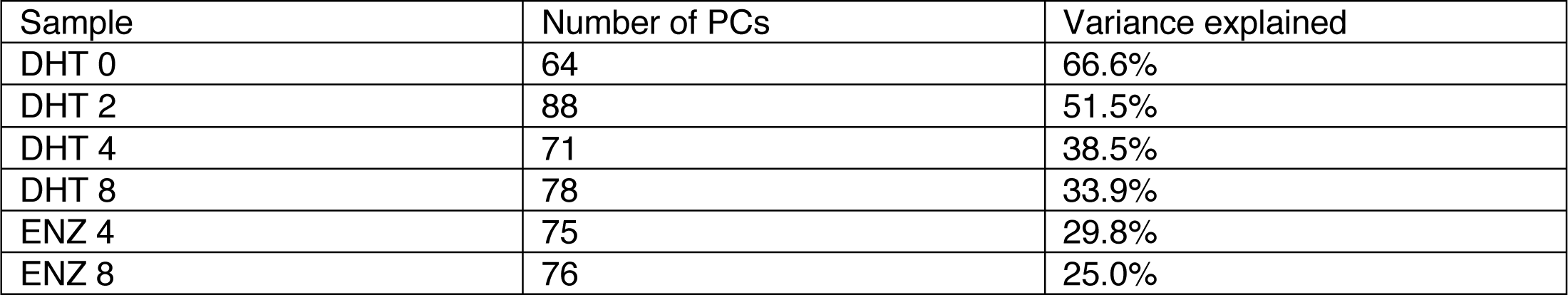

| Sample | Number of PCs | Variance explained |
| --- | --- | --- |
| DHT 0 | 64 | 66.6% |
| DHT 2 | 88 | 51.5% |
| DHT 4 | 71 | 38.5% |
| DHT 8 | 78 | 33.9% |
| ENZ 4 | 75 | 29.8% |
| ENZ 8 | 76 | 25.0% |

Using these PC loadings, we construct an FDL for each sample for visualization using the draw graph function in scanpy(*15*). Each FDL is based on the affinity graph of each sample (kNN=30; kernel width σ = 10).

#### SEACell algorithm for calculating metacells

A key challenge in scRNA-seq data analysis is that much of the observed variability across cells is due to limitations in molecular sampling; each profile only quantifies a small fraction of the total transcripts in each cell. Clustering of cells can help overcome this sparsity by averaging out noise, but this strategy fails to capture the full heterogeneity of cell states in the dataset. To retain a complete description of phenotypic diversity while simultaneously mitigating noise and sparsity, we aim to aggregate cells that share the same cell state at a much finer resolution than clustering.

Our strategy is to group cells into metacells. Each metacell represents a distinct, highly granular cell state, whereby differences between cells within a given metacell are due to technical rather than true biological differences. Critically, single cells within each metacell should follow an identical statistical distribution of gene counts. Aggregating counts across all cells in each grouping produces a robust, full transcriptomic quantification of that cell state, which overcomes noise and gene dropout. Moreover, metacells enumerate the observed cell states in the data and enable downstream computation on these states in a manner that is unbiased by their frequencies.

The concept of metacells was introduced in Baran et al.(*16*), but the accompanying method was developed for studying healthy cells and it removes outliers aggressively. In contrast, tumors typically exhibit heterogeneity that is rife with outliers of biological significance, as we have also observed in our study of plasticity following *Trp53* and *RB1* deletion.

We therefore used our recently developed “Single-cEll Aggregation for high resolution Cell states” (SEACell) method, which is better suited to our data. SEACell requires few parameters, is robust to parameter settings, is able to handle outliers, and takes advantage of graph-based manifold learning algorithms that have proven to capture the landscape of cell states in single-cell genomics data faithfully and robustly. A full treatment of the SEACell algorithm and its benchmarking will appear in Persad et al. (manuscript under preparation), but here we include sufficient detail about the algorithm and its performance for our application.

As input, the SEACell algorithm accepts a count matrix of individual cells and associated PC loadings. Here, we normalize the UMI counts per sample *U_s_* by library size, scale by median library size, and perform natural log transformation with pseudocount of 1 for subsequent aggregation of counts per metacell. A k-nearest neighbor (kNN) graph is constructed using Euclidean distance between cells in PC space. To remove spurious edges and obtain a more faithful representation of the phenotypic manifold, we construct a shared nearest neighbor (SNN) graph: we prune edges in the kNN graph that are not bidirectional and weight remaining edges using an adaptive Gaussian kernel following van Dijk et al.(*14*), which accounts for variation in density along the manifold.

SEACell then partitions the SNN graph into cell groupings using archetypal analysis, a method to fit complex phenotypic data to a lower-dimensional convex polytope amenable to study(*14,17*). Here, archetypes represent points that define extrema in the SNN graph, such that observations can be approximated by convex combinations of the archetypes. This framework naturally yields a soft assignment of each single cell to the selected SNN archetypes, forming metacells.

Archetypal analysis proceeds as follows. Let matrix *X* ∈ ℝ*^c^*^×*c*^ represent the adjacency matrix of the SNN graph. Archetypal analysis aims to identify *m* archetypes by computing matrices *A* ∈ ℝ*^c^*^×^*^m^* and *B* ∈ ℝ*^m×c^* that minimize the objective function

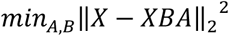

Subject to the following constraints:

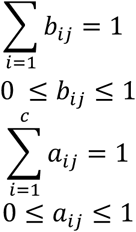

*B* represents the archetype matrix that defines archetypes as linear combinations of columns in *X*, whereas *A* is an assignment matrix that assigns each cell to one of the archetypes specified by *B*. In this way, each cell in *A* is an affine combination of weights for the selected archetypal cells, yielding a soft assignment of cells into archetypes. We can also perform a hard assignment of each cell to an archetype based on the highest weight. These final archetypes represent our set of SEACell metacells.

Solving the above objective function proceeds by gradient descent. Archetypes are first initialized by greedily selecting columns in ( that maximally reduce the Frobenius norm between *X* and the projection of *X* onto those columns(*18*). This greedy approach iteratively selects each column to maximize the remaining explained variance, such that the remaining linearly independent combination of cells is well distributed over the phenotypic manifold.

The number of metacells *X* is a parameter that needs to be set for each sample analyzed. The optimal value can be chosen using robustness analysis—by generating metacells over a large range of *X*, computing the adjusted Rand index (ARI) between all pairs of metacell datasets, and identifying a regime of *m* values with high ARI, representing robust metacell groupings (see section below).

The output of SEACell includes per-cell weights for each metacell, per-cell hard assignment to each metacell, and the single cell closest to the centroid of each metacell. The algorithm aggregates counts within each metacell. Aggregated counts may then be normalized and scaled by median library size, and log-transformed for downstream applications.

#### Identifying robust metacells in the organoid time course

To characterize the wildtype phenotypic landscape and how it remodels following perturbation, we used SEACell to calculate metacells separately within each sample timepoint *t*.

We first determined the number of metacells for each timepoint *m_t_*by calculating metacells over the range [20, 500] in increments of 10 (**Fig S4A**). We chose the kneepoint in the distribution of ARI between consecutive values of *m* as the final *m* for each timepoint (**Fig S4B**). In this way, we choose a minimum *m_t_* that is robust based on ARI, with a median metacell size of 15.5 across timepoints. Based on this final selection of *m* for each sample, the median size of each metacell grouping ranges from [12, 24] across timepoints (**Fig S4C**). The centroids of each metacell grouping are well distributed over the corresponding FDL of single cells individually constructed for each sample (**Fig S4D**) as well as across all samples combined (**Fig S4E**).

Next, we aggregated counts within each metacell. We normalized and scaled the aggregated counts by median library size and performed natural log transformation with a pseudocount of 1. We selected the 5000 most variable genes for PCA and retained the top 50 PCs, which explain 82.0% (DHT 0), 77.5% (DHT 2), 70.3% (DHT 4), 80.7% (DHT 8), 70.3% (ENZ 4), and 77.9% (ENZ 8) of the variance. The variance explained in each sample by the top 50 PCs is much higher for metacells than for single cells, demonstrating the greater fidelity and robustness provided by SEACell aggregation. For each sample, we constructed a kNN graph for visualization using a force-directed layout. To ensure that graph connectedness scales proportionately with number of metacells, we set kNN to the ceiling of 3% of *m* (kNN = 6 in DHT 0; kNN = 5 in DHT 2; kNN = 6 in DHT 4; kNN = 4 in DHT 8; kNN = 6 in ENZ 4; kNN = 4 in ENZ 8).

To assess the combined dataset, we pooled metacells identified from each individual timepoint, normalized as for individual timepoints, then selected the 5000 most variable genes for PCA and retained the top 50 PCs, which explain 68.3% variance. We can visualize these metacells using a kNN graph (kNN = 10) with an FDL.

**Figure S5** shows a flow chart of data-processing, highlighting what analysis was applied directly on the single-cell data and what analysis was performed at the level of metacells.

#### Identifying basal and luminal subtypes in the wildtype organoid

To assess changes following *RB1/TP53* deletion, we first needed to characterize basal and luminal subpopulations in the wildtype organoid. We grouped individual cells from the wildtype into SEACell metacells and performed unsupervised clustering using Phenograph(*19*) with kNN = 7 (4% of the number of metacells, as above), which identified 5 clusters. We aimed to characterize these 5 wildtype subpopulations by 1) correlating to sorted, bulk-sequenced basal and luminal cells, 2) calculating differential expression, and 3) scoring key gene sets.

First, we correlated each metacell to our bulk RNA-seq reference dataset of FACS-sorted CD49f+ basal and CD24+ luminal cells (2 replicates each; see **Bulk RNA sequencing and analysis**). We pre-processed the bulk dataset by averaging log2(TPM) across replicates, yielding one basal cell and one luminal cell profile. To facilitate comparison, we z-scored gene expression across observations in the metacell dataset as well as across the bulk RNA-seq dataset. To ensure robust discrimination between basal and luminal cell types, we performed feature selection *F* on significant differentially expressed genes (DEGs) in each wildtype cluster compared to all other clusters (abs(coef) > 1 and Bonferroni-adjusted p-value < 0.01; see **Differential expression**) and excluded genes from cell cycle, hypoxia, and apoptosis pathways that are not specific to cell type and might confound classification. The filtered genes included pathways REACTOME_CELL_CYCLE_MITOTIC, REACTOME_MITOTIC_G1_G1_S_PHASES, HALLMARK _G2M_CHECKPOINT, HALLMARK_HYPOXIA and HALLMARK_APOPTOSIS from MSigDB as well as a list of cell cycle genes “regev_lab_cell_cycle_genes.txt” from the scanpy package(*15*). This filter *F* left 961 cell-type-specific DEGs (Table S8), which we used for calculating Pearson’s correlation to the bulk basal (or luminal) profile. Since our initial z-score transformation centers bulk CD49f+ and CD24+ dataset gene expression at 0, high correlation to the basal cell line corresponds to low correlation to the luminal cell line, with near symmetry at 0. To avoid redundancy, we therefore consider correlation to basal cells alone for subsequent analysis. We find that each metacell cluster generates correlations significantly different from a null hypothesis of 0 (Bonferroni-corrected one-sample t-test), rendering each cluster readily classifiable as either basal or luminal (**Fig S6B**). We therefore identify 3 wildtype basal subpopulations (Basal Org1, Basal Org2, Basal Org3) and 2 wildtype luminal subpopulations (Luminal Org1, Luminal Org2).

Close inspection of known cell type markers among DEGs for each wildtype cluster supports these basal versus luminal designations. A select subset of these DEGs are shown in **Figure 2C**. Basal subpopulations uniformly express canonical basal markers (*Krt5, Krt14, Trp63, Itga6*), and luminal subpopulations express luminal markers (*Ly6d, Cd24a*), although Luminal Org1 expresses additional luminal markers (*Krt4, Krt6a, Dpp4*) with some weakly expressed basal features (*Krt5, Krt14, Trp63*).

#### Assessing basal-luminal mixing following *Rb1/Trp53* deletion

We next considered the problem of assigning basal or luminal cell types to the mutant organoid, which represents a highly plastic system that has shifted from its original phenotypic identity. We followed several orthogonal strategies to quantify plasticity as a vehicle of basal-luminal mixing: 1) Markov absorption in single cells, 2) gene set scores in single cells, and 3) diffusion distances in SEACell metacells. Each of these strategies has different advantages and drawbacks, but all consistently model increased basal-luminal mixing over time following *Rb1/Trp53* loss and with ENZ treatment.

##### Modeling plasticity using Markov absorption classification of single cells

Although SEACell allows more robust quantification of gene expression for each state in a sample (see **SEACell algorithm for calculating metacells**), we initially sought a method that can deconvolve basal-luminal mixing at the single-cell level. We consider the following semi-supervised classification of unassigned cells to a mixture of 5 basal and luminal subtypes from the wildtype organoid. For *N* cells, where a subset of *L* cells has known phenotype (training data), we must assign the remaining *N-L* cells (test data) the probability of representing subtype *S ϵ {s_1_,s_2_,s_3_,s_4_,s_5_} = {*Basal Org1, Basal Org2, Basal Org3, Luminal Org1, Luminal Org2*}*.

To assess basal-luminal mixing using phenotypic similarities between cells, we represent the dataset as an absorbing Markov chain and follow the graph-based classification approach first demonstrated in Levine et al.(*20*). The Markov chain is represented as a kNN-graph, where each node is a cell belonging to the set of unassigned cells *C = {c_1_,…,c_N_}*. We also consider the set *Y = {y_1_,…,y_N_}* of subtype assignments where *y* ∈ *S* and *S* is the set of all possible subtype assignments. Each edge corresponds to the transcriptional similarity between cells. For cell *c*, we want to calculate the probability of its assignment to a particular subtype *s*, or *Pr(yc == s)*. Given the subtype labels of *Y = {y_1_,…,y_L_}*, we must assign subtype labels to the remaining unassigned cells or test data *Y_L+1:N_ = {y_L+1_,…y_N_}*. We treat the labeled cells as an absorbing state where a random walk begins from an unassigned cell and ends when it reaches a classified labeled cell. Therefore, *Pr(yc == s)* is reformulated as the probability that a random walk from *x* will reach a labeled node of subtype *s* first. We compute this for all possible random walks that end in all labeled nodes of subtype *s* using an analytical solution for calculating these probabilities, which is implemented in the Phenograph package(*19*).

To construct the Markov graph, we consider the combined dataset where our wildtype cells serve as labeled training data. We again use the selected features *F* (including wildtype DEGs and excluding genes related to cell cycle, hypoxia, and apoptosis) (Table S8) and project log-normalized counts at the single-cell level (see above) onto the first 65 PCs selected by knee-point detection, corresponding to 82.8% variance explained. We then constructed a kNN-graph with k=25 and obtain the first 18 diffusion components (DCs) retained by eigengap. Using the Phenograph package, we transform this diffusion graph into a Jaccard graph between k-neighborhoods, which has been shown to be more robust to noise. The resulting graph represents a Markov chain, which can be used to calculate the Markov absorption probabilities for each unlabeled cell *c* to reach a labeled cell of a given subtype *s*. We next consider the *per-*cell Markov absorption probabilities *M_cs_* of each basal or luminal subtype *s*, where ∑*_s∈s_ M_cs_* = 1, representing a deconvolution of mixed phenotypes per cell. We can then use the entropy of subtype probabilities per cell *H_c_* as a measure of basal-luminal mixing, or plasticity, where

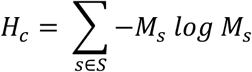

(**Fig 2F**). The *per*-cell entropy scores are remarkably robust to the number of DCs retained from 15 to 24 (**Fig S6G**).

To better visualize the mixed phenotypes per cell, we consider that three of the wildtype organoid subtypes (Basal Org2, Basal Org3, Luminal Org2) are predominantly represented in mutant cells. We can readily represent the deconvolution of these three phenotypes in a 3-coordinate ternary graph, as implemented in the ggtern package(*21*)(**Fig 2E**).

##### Modeling plasticity using gene set scores in single cells

To confirm our finding of basal-luminal mixing, we consider an orthogonal approach that is independent of initial kNN graph construction based on cell-cell similarity, instead only considering the DEGs computed for each wildtype subpopulation. For each cell *c* in the organoid time course, we calculate the *per*-cell subtype score *Z_cs_* as the average z-score of DEGs for each wildtype basal/luminal subpopulation *s*. We normalize *Z_cs_* such that ∑*s∈S Z_cs_* = 1. We can again use entropy of subtype scores per cell *H_c_* as a measure of basal-luminal mixing, or plasticity, where

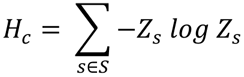

(**Fig S6I**). To better visualize deconvolution, we can consider the three predominant subtypes (Basal Org2, Basal Org3, Luminal Org2) and plot these cells on ternary plots, as above (**Fig S6H**).

##### Modeling plasticity using diffusion distance in SEACell

We can represent the metacell phenotypic manifold in the same manner as for single cells, by constructing a kNN graph and visualizing with a force-directed layout. In our rendering of the FDL, we colored each SEACell metacell based on its correlation to the reference CD49f+ profile. We observed increasing interconnections between basal-like and luminal-like metacells across time and treatment. Many previous approaches treat paths along edges of the kNN-graph as potential paths for cell state transitions(*22-24*), though the direction of transition is unclear in this application. Nevertheless, because our SEACell formulation creates a set of distinct cell states, the increasing interconnections are unlikely to arise between densely sampled cells of the same cell state. We therefore surmised that increasing interconnections could indicate increasing routes of transdifferentiation between basal and luminal states and thus serve as a good proxy for plasticity.

We can formalize this observation using a two-step strategy based on 1) attributed stochastic block models (aSBM) and 2) diffusion maps. First, let *G* be an undirected kNN-graph of metacells. Each metacell carries a node attribute based on the correlation to bulk-sorted CD49f+ basal versus CD24+ luminal cells (see **Identifying basal and luminal subtypes in the wildtype organoid**). We seek to partition *G* into a connected basal subgraph *GB* and connected luminal subgraph *G_L_*, so that we can later assess the connectivity between *G_B_* and *G_L_*. One popular approach for community detection is the stochastic block model (SBM), which is a generative model that probabilistically creates edges within and between communities. The number of communities *k* is pre-specified by the user. This model can be fitted to the observed graph so that nodes are partitioned into communities that maximize the likelihood of the model. The traditional SBM determines community membership through connectivity alone; however, we wish to partition graphs based on the node attribute of basal versus luminal correlation as well. We therefore consider the attributed stochastic block model (aSBM)(*25*), which estimates community assignment by simultaneously considering the adjacency matrix *A* (connectivity) and a node vector of continuous attributes *N* (basal vs luminal correlation). *A* and *N* follow Gaussian mixture models with parameters that are each assumed to be conditionally independent given community membership. We follow an approach by Stanley et al., which uses expectation-maximization (EM) to maximize a likelihood function that estimates both community membership and parameters of the Gaussian mixture model. Applied to our organoid data, we use aSBM to partition *G* into *k* = 2 basal and luminal communities *G_B_* and *G_L_*. We initialize our aSBM using hard classification based on basal versus luminal correlation. EM then converges upon a solution of *G_B_* and *G_L_* that maximizes the likelihood function based on both connectivity (within versus between partitions) and the difference between node attributes within each partition. The final result is the identification of two connected components corresponding to predominantly basal or luminal cells.

Next, having identified basal and luminal compartments *G_B_* and *G_L_* for each graph, we seek a summary statistic that captures the increasing interconnections between compartments *G_B_* and *G_L_* as a proxy measure of plasticity. For a randomly selected basal cell *B* in *G_B_* and *L* in luminal cell in *G_L_*, we consider the diffusion distance between them, a popular metric to measure distance along the phenotypic manifold in single-cell data analysis(*22, 23*). The diffusion distance between *B* and *L* reflects the connectivity between two points, considering the probability over all paths in the graph. Assuming the neighbor-graph represents potential paths of cell-state transitions between similar cell states, the diffusion distance quantifies the potential to transdifferentiate between basal and luminal states, where a smaller distance corresponds to a greater potential for transdifferentiation.

Mathematically, we construct the diffusion map *E* ∈ ℝ*^c^*^×*L*^, where A is the number of the top diffusion components (DCs), corresponding to eigenvalues λ_1_, λ_2_, …, λ_L_ of the adjacency matrix repressing the kNN-graph. Diffusion distance *DD* between cells, *i* and *j* can be calculated as:

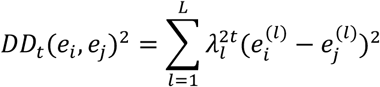

where *t* is the number of steps through the graph and *e_i_^(l)^* is the embedding of cell *i*, along diffusion component *l*. To avoid setting *i*, we calculate a multi-scale diffusion distance *MS* following Setty et al.(*23*) that accounts for all scales at once:

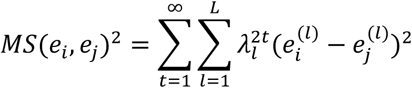

Because 1 > λ_1_ > λ_2_ > … > λ_1_, the above equation can be rewritten as

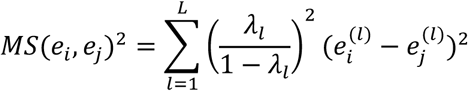

This multi-scale diffusion distance is robust to the number of diffusion components retained, outlier cells, and density differences along phenotypic space.

To apply this approach to the combined metacell dataset, we construct the diffusion map based on the top 50 PCs as described in **Identifying metacells in the organoid timecourse** and retain the top 6 DCs based on eigengap. For each sample, we then consider the log mean *MS* between pairs of basal and luminal metacells as an inverse measure of plasticity. To compare *MS* across sample timepoints and control for differing numbers of metacells in each timepoint, we randomly subsample 50 basal and 50 luminal metacells for 1000 iterations, each time calculating the distribution of pairwise diffusion distances and taking the log10 mean. We found that the log mean *MS* is robust to selection of diffusion components from 5 to 15, based on Pearson’s correlation (**Fig S6J**).

#### Robustness analysis

We ensured our analysis was robust in several ways. With respect to robust metacell identification, please see **Identifying robust metacells in the organoid timecourse**. With respect to our *per*-cell entropy measure for plasticity based on cell type classification using Markov absorption (please see **Modeling plasticity using Markov absorption classification of single cells**), we repeated our entropy calculation over a number of different DCs retained from 15 to 24. We measured the Pearson’s correlation between entropy measures and found that all pairwise correlations are well above 0.9, ensuring robustness (**Fig. S6G**). With respect to our plasticity score based on the log mean multi-scale diffusion distance between metacells (please see **Modeling plasticity using diffusion distance in SEACell**), we repeated calculations over a number of different DCs retained from 5 to 15. We again measured pairwise Pearson’s correlation between multi-scale diffusion distances based on different DCs and found that correlations are well above 0.9, ensuring robustness (**Fig S6J**).

#### Differential expression

We performed differential expression comparisons between 1) mutant and wildtype metacells in the combined timecourse, and 2) each metacell cluster and all other clusters in the wildtype condition. All differential expression was performed using MAST (version 1.8.2)(*26*), which provides a flexible framework for fitting a hierarchical generalized linear model to the expression data. We used a regression model that adjusts for cellular detection rate (cngeneson, or number of genes detected per sample):

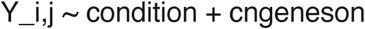

where condition represents the condition of interest and *Y_i_* is the normalized expression level of gene *i* in cells in cluster j, transformed by natural logarithm with a pseudocount of 1. We considered genes to be significantly differentially expressed for Bonferroni-adjusted p-value < 0.1 and absolute log fold-change > 1.

#### Identifying gene signatures in single-cell data

Enriched gene pathways were identified using pre-ranked GSEA, as implemented by the R package fGSEA(*27*), based on a pre-ranked gene list using 10,000 permutations. Gene ranks were calculated using -log(p-value)*logFC based on MAST(*26*) differential expression (described above). To assess enriched pathways, we used a curated set of pathways from MSigDB v 7.1(*28*)(*29*) (**Data S1**). Using the same cutoff as in the original GSEA paper, we considered pathways with Benjamini-Hochberg adjusted p-values < 0.1 to be significant, as well as abs(NES) > 1, where NES is the normalized enrichment score. To score activation of significantly enriched pathways in our dataset, we would consider the leading-edge gene subset based on GSEA and calculate the average z-score of the gene set.

#### Assessing Putative Pathways Driving Plasticity

We sought to identify candidate pathways that may lead to increased plasticity in the mutant organoid. We reasoned that such gene programs would be 1) correlated to our measure of plasticity and 2) expressed early in the time course (2^nd^ week), when plasticity begins to rise.

To this end, we first assessed the correlation of pathways with plasticity for each single cell. We considered the entropy of Markov absorption cell-type probabilities as a measure of plasticity (see **Modeling plasticity using Markov absorption classification of single cells**) and used the set of curated pathways assessed by GSEA (see **Identifying gene signatures in single-cell data**) to calculate pathway scores. Each pathway was scored by the average z-scored expression of the gene set. For significant pathways defined by abs(NES) > 1 and Benjamini-Hochberg-adjusted p-value < 0.01, we considered the leading-edge gene subset, which is the core set of genes in a given pathway that drive enrichment(*29*). We then calculated the Pearson’s correlation between plasticity and each pathway score (**Table S11**).

Beyond the association between plasticity and pathway score, we reasoned that timing would be an indication of possible causality, given that participating gene programs should be expressed prior to the appearance or expansion of plasticity. Among the top 10% of pathways correlated with plasticity, we ordered gene programs that increase early in the time course (week 2). We show select pathways from the top 10% in **Figure 3A** filtered for pathway redundancy and identify LIF and JAK/STAT, IFN alpha/beta, and FGFR signaling as possible drivers of plasticity.

#### Cell-cell interaction analysis

A number of methods for inferring ligand-receptor (L-R) interactions have been developed for single-cell analysis. Most focus on identifying paracrine interactions, whereby a ligand expressed by a sender cell binds to a receptor expressed by a receiver cell. The most common strategies assess L-R differential expression across cell types (i.e. CellTalker (*30*) or L-R co-expression in comparison to a permuted null model (i.e. CellPhoneDB (*31*). A critical caveat of these methods is that they make strong assumptions on interactions between each pair of sender and receiver cell. This concern is mitigated in our framework, given our focus on autocrine signaling sets the sender and receiver to be the same cell.

An additional caveat of most current strategies is that they disregard downstream signaling in the receiver cell, which is key to distinguishing functional interactions from non-interacting L-R pairs that happen to be co-expressed. NicheNet(*32*) is best known among the very few methods that incorporate knowledge of downstream signaling. It prioritizes L-R candidates by calculating ligand-target regulatory potential; however, potential is based solely on a static network of ligands, transcriptional regulators and target genes mined from the literature, and it fails to incorporate the actual expression of transcriptional regulators or target genes in the dataset. Moreover, the default model of downstream signaling is constructed from multiple literature sources that mix diverse biological contexts and is therefore unlikely to be representative of any single cell type. It is left to the user to curate and apply to specific datasets. This static model is blind to signaling crosstalk that is likely to occur in any given system.

We were explicitly interested in identifying ligand-receptor interactions that could explain the JAK-STAT signaling observed in the mutant organoid cell line. We benefited from the study of our simple organoid model, such that we only needed to consider autocrine L-R interactions and were able to exclude cell type as a variable in L-R analysis. We therefore sought to reverse-engineer the problem of cell-cell interactions to consider only autocrine L-R candidates that could explain the observed increase in JAK-STAT signaling. To restrict our search space, we considered a curated set of 74 L-R pairs known to activate JAK-STAT signaling (**Table S12**).

We then sought a *per*-cell summary statistic of JAK-STAT signaling. To this end, we first obtained a gene signature for observed JAK-STAT signaling specific to our dataset by combining the leading-edge subsets of KEGG_JAK_STAT_SIGNALING_PATHWAY and HALLMARK_IL6_JAK_STAT3_SIGNALING as identified by GSEA, leaving a set of 33 genes (Table S13). We then defined the JAK-STAT signaling score *JS* as the average z-score of these 27 genes. For each pair of ligand expression *L* and receptor expression *R*, we then performed linear regression over all cells using the formula *JS ∼ L + R + L: R.* We scored how well each *L-R* pair explains *JS* using the adjusted Radj^2^. Considering non-zero Radj^2^ narrowed the list to 15 possible L-R candidates (**Fig 4A, S8A**).

In addition to how well the L-R expression fits JAK/STAT signaling, we prioritized candidate drivers of JAK/STAT signaling and subsequent plasticity using two more criteria: receptor expression *R* that is upregulated 1) early in the timecourse (week 2), and 2) with treatment. These additional criteria are profiled in **Figs. 4B, S8B,** and **S8C,** and prioritized LIF (co)receptors and FGF receptors as plausible drivers

